# Persistent transcriptional programs are associated with remote memory in diverse cells of the medial prefrontal cortex

**DOI:** 10.1101/784413

**Authors:** Michelle B. Chen, Xian Jiang, Stephen R. Quake, Thomas C. Südhof

## Abstract

It is thought that memory is stored in ‘engrams’, a subset of neurons that undergo learning-induced alterations. The role of gene-expression during learning and short-term memory has been studied extensively, but little is known about remote memory that can persist for a lifetime. Using long-term contextual fear memory as a paradigm, an activity-dependent transgenic model for engram-specific labeling, and single-cell transcriptomics we probed the gene-expression landscape underlying remote memory consolidation and recall in the medial prefrontal cortex. Remarkably, we find sustained activity-specific transcriptional alterations in diverse populations of neurons that persist even weeks after fear-learning and are distinct from those previously identified in learning. Out of a vast plasticity-coding space, we uncover select membrane-fusion genes that could play important roles in maintaining remote memory traces. Unexpectedly, astrocytes and microglia also acquire new persistent gene signatures upon recall of remote memory, suggesting that they actively contribute to memory circuits. Our discovery of novel distinct gene-expression programs involved in long term memory adds an important dimension of activity-dependent cellular states to existing brain taxonomy atlases and sheds light on the elusive mechanisms of remote memory storage.

---

Memory is the function of the brain by which information is encoded, stored, and retrieved, and is critical for adaptation and survival. Long-term memories do not form immediately after learning, but develop over time through a process of stabilization, known as consolidation ^1–3^. Previous studies have uncovered important roles of a series of molecular and cellular processes in learning and memory, such as CREB-dependent gene expression, cAMP signaling, and synaptic plasticity ^4–8^. The central role of RNA synthesis and subsequent protein translation was first shown in mice, and mechanistically studied in many organisms^9^. Despite these discoveries, the molecular underpinnings of memory consolidation remain elusive. For instance, while gene expression alterations are found in the first 24 h of learning, it is not clear whether these changes are sustained or new ones are acquired in order to consolidate a long-term memory trace that is resistant to disruption on the scale of weeks or years ^10–12^. Moreover, the dependence of long term memory on the hippocampus is known to degrade over time, with cortical structures being increasingly important^13^. Recently, the development of transgenic tools that allow the identification of small activated neuronal ensembles based on immediate-early-gene (IEG) expression have allowed the dissection of the molecular mechanisms underlying experience-dependent connectivity and plasticity ^14–16^. Here, we combine a system for activity-dependent genetic labeling (TRAP2) to access engram-specific cells ^17–19^, an established model of learning and memory (contextual-fear conditioning), and single-cell transcriptomics to uncover the role of gene expression in long term memory storage. We find that cells activated by remote memory recall exhibit sustained transcriptional changes that are both activity-dependent and experience-specific, and that are distinct from the genes expressed during memory encoding. Importantly, we begin to resolve the heterogeneity of plasticity mechanisms via identification of specific genes that have the potential to regulate, enhance, or disrupt long term memory storage.

## Molecular characterization of active neuronal populations during remote memory consolidation

The ability to resolve and characterize an experience-specific neuronal ensemble out of a vast background of neurons is crucial in our understanding of the molecular code that regulates memory formation and storage. The prevailing model for neuronal representation of memory is the Engram Theory ^20^, which posits that learning activates a small ensemble of connected neurons in the brain and induces persistent physical and biochemical changes in the connections between these neurons. These so-called engram cells or “trace cells”, provide physical locations for the storage and retrieval of memories ^21,22^. *TRAP2;Ai14*, a new cFosTRAP transgenic mouse line, is a system that enables the identification of activated neurons via fluorescence by using the immediate-early gene (IEG) *c-Fos* locus to drive the expression of tamoxifen inducible CreER, along with a permanent Cre-dependent tdTomoto reporter ^19^. To identify the transcriptional programs promoting memory consolidation in contextual fear learning, we trained *TRAP2; Ai14* mice in a conditioning chamber with 3 pairs of tone-foot shocks and returned the mice 16 days later for remote contextual memory recall. 4-OHT was injected into the animals immediately prior to memory recall in order to label experience-specific neuronal ensembles. The fear-conditioning followed by recall training paradigm (fear-recall: FR) identifies IEG-expressing neurons during the recall event, the ensemble where memory has been consolidated ^19,23^. In addition, three control groups were implemented: return to context without fear conditioning (no-fear: NF), fear-conditioning without recall (no-recall: NR), and no fear conditioning-nor return (homecage: HC) (**Fig 1a-c**). Together, the collection of TRAPed neurons from each training group allows the comparison of memory consolidation-specific activation versus various forms of baseline activity.

**Figure 1.**
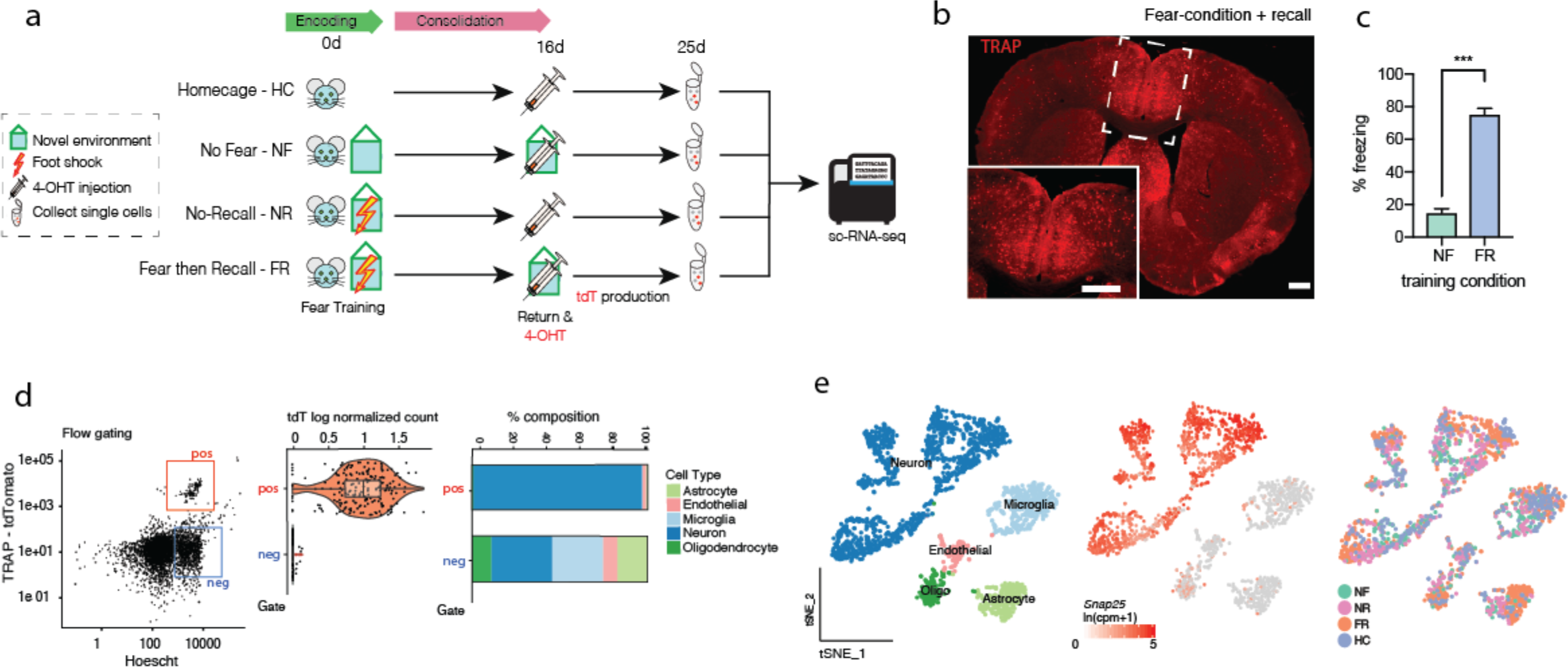
Labeling and collection of single neurons in activated memory traces via c-Fos TRAP. a) Experimental paradigm of generating remote fear-memory traces via contextual fear conditioning, isolation of TRAP+ activated neurons via flow cytometry, and subsequent full-length plate-based sequencing. b) Representative image of TRAPed tdT+ (red) cells 9d after 4-OHT injection during remote recall. c) Degree of freezing upon return to the novel context 16 d after fear conditioning (FR) or no conditioning (NF) (n=4 mice per condition). d) Representative flow gating for *tdT*+ TRAPed cells. Post-sequencing analysis shows enrichment of tdT mRNA in the positive sort gate, an enrichment of neuronal cell types (*Snap25*+) in the positive gate, and a distribution of neuronal and non-neuronal cell types in the negative gate (Extended Fig 1b). e) tSNE reduction of all sorted cells with neuronal cells identified via expression of *Snap25* mRNA. All training and control conditions are represented in all cell clusters.

Samples were collected from the medial prefrontal cortex (mPFC), a cortical region heavily implicated in remote but not short-term memory ^24,25^. To determine whether persisting transcriptional changes exist in the mPFC and to identify such changes, we performed deep plate-based single-cell mRNA sequencing on both TRAP+ and TRAP− cells enriched via flow cytometry, 9 days after the retrieval event (25 days after fear-learning) (**Fig 1d**). This time frame allows adequate time for the *tdT* reporter protein to be expressed ^18^, and gives access to the persistent transcriptional state of recall-activated cells resulting from remote memory consolidation. The percentage of TRAPed cells collected was significantly higher in FR (~1.5% of all cells) than in other conditions (**Extended Fig 1a**), confirming that the TRAP2 activation captured increased neuronal activity during the fear-recall process. In total, we sequenced 3691 neuronal cells (Snap25+/tdT+ or tdT−) and 2672 non-neuronal cells from all 4 behavioral trainings, with an average read depth of 1 million reads/cell. An average of ~6000 expressed genes were identified in *Snap25*+ neurons (**Extended Fig 1b-c**). Unbiased transcriptome clustering of all cells from all four training conditions allowed the identification of major cell types and confirmed the dominance of neurons in the positive sorting gate, whereas both non-neuronal and neuronal cells (*Cldn5*+ endothelial, *Pdgfra*+ OPCs, *Cx3cr1*+ microglia, *Aqp4*+ astrocytes) were present in the negative gate (**Fig 1d-e**, **Extended Figure 1d)**. The enrichment of *tdTomato* mRNA in our putative positive gate was also confirmed. Both TRAP+ and TRAP− cells from FR training groups and three controls (NR, NF, and HC) were represented in all clusters, suggesting that neither the neuronal activation state nor the training paradigm significantly alters cell-type identities on a larger scale (**Fig 1d**).

Sub-setting and re-clustering all 3691 neurons revealed 7 putative neuron sub-populations - 4 glutamatergic (C0, C2, C3, C5) and 3 GABAergic (C2, C4, C6) - all of which were consistently observed throughout 4 biological replicates (**Fig 2a-c, Extended Fig 2a**). Analysis of enriched genes per neuronal subtype show molecularly distinct populations (**Fig 2d**), with each subtype expressing at least one highly distinctive marker (>90% expression, **Fig 2e**). All subtypes contain *tdT*+ cells, indicating the ability of all neuron subtypes to be activated, regardless of the training states (**Extended Fig 2b**). Comparisons of key marker genes (C0-*Dkkl1,* C1-*Rprm*, C2-*Calb2*, C3-*Tesc*, C4-*Tnfaip8l3*, C5-*Tshz*, C6-*Lhx5*) to existing cortical single cell expression databases (Zeisel et al., 2018; Allen Brain Atlas) confirmed their presence in the mPFC as well as their layer specificities (**Extended Fig 2c**).

**Figure 2.**
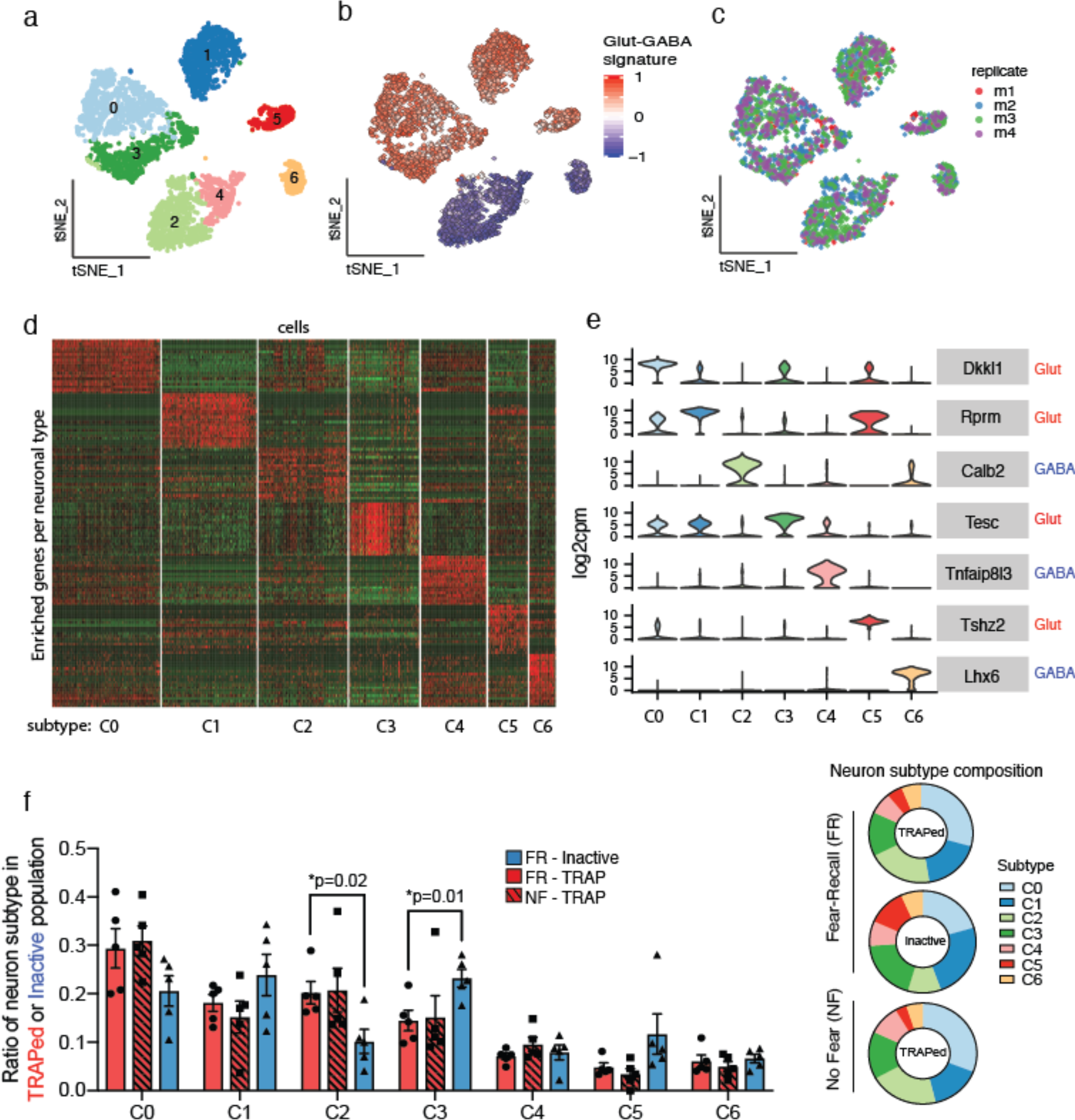
Molecular identification of activated neurons during consolidation in a remote memory trace. a) tSNE reduction and Louvain clustering of all *Snap25*+ neurons (total 3691 cells across 4 conditions) using the top 2500 variable features reveals 7 distinct neuronal subtype (C0-C6). b) Identification of excitatory (glutamatergic) and inhibitory (GABAergic). Glut-GABA signature is calculated based on the difference of the scaled expression level of *Gad1* and Slc17a7. c) Distribution of biological replicates between C0 to C6 subtypes for all training conditions combined. d) Heatmap depicting the top 25 enriched genes per neuron subtype and distinctiveness of their gene expression. e) Violin plots of expression of putative marker genes for each neuron subtype (C0 to C6) (from d). f) Bar plot of the differences in neuron subtype composition of TRAPed populations in FR and NF mice, as well as inactive populations in FR mice (n=5 mice/condition). Ring plots depict the distribution of neuronal types (C0-C6) in each group of cells collected.

To understand whether fear memory consolidation involves the differential activation of distinct neuron subtypes compared to baseline activity, we compared the subtype composition of TRAPed populations collected from FR and NF control mice (**Fig 2f, Extended Fig 2d**). Surprisingly, no significant differences between the FR and NF groups were found in the ratios of the 7 subtypes. This suggests a lack of training-dependent recruitment of neuron types during consolidation compared to basally active populations in a no-fear memory scenario. Excitatory and inhibitory neuron types were both represented in active FR populations, with glutamatergic cells composing ~60-70%. Additionally, within the same FR brains, active and inactive populations had roughly similar neuron subtype compositions, with the exception of C2-*Calb2* and C3-*Tesc*, suggesting only slight shifts in the recruitment or retirement of neuron subtypes due to activity.

## Memory consolidation activates long-lasting transcriptional programs that are heterogeneous across neuron subtypes

To determine whether consolidation-dependent transcriptional changes occur, we looked for differentially expressed genes (DEGs, log2FC>0.3 and FDR<0.05) in TRAPed FR vs NF cells (**Fig 3a**). Single cell resolution enables the comparison of neurons within the same subtype, and comparing active populations allows the discovery of transcriptional programs that are related to memory consolidation rather than baseline activity (**Extended Fig 3a**). Of 23,355 genes, 1292 were found to be consolidation-dependent, and expression patterns indicated an overall transcriptional activation, with more genes up- than down-regulated. Interestingly, DEGs were largely distinct between subtypes (**Fig 3b**). This subtype-dependent heterogeneity could be indicative of specialized functional roles rather than one mass population working in unison within the memory trace.

**Figure 3.**
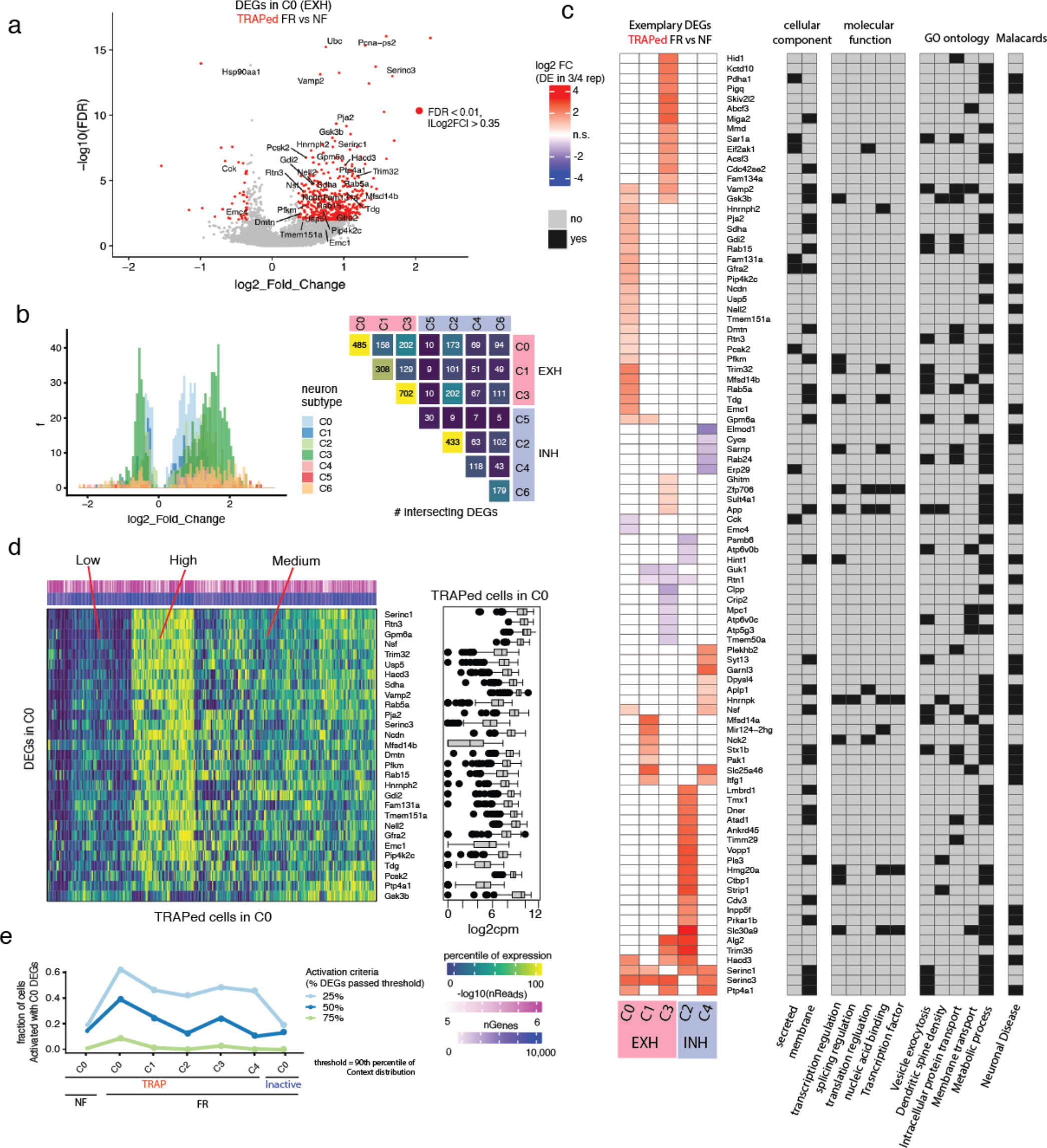
Transcriptional programs activated during consolidation of remote memories are distinct per neuron subtype. a) Volcano plot for neuron type C0 (n=126 (NF) and 289 (FR) cells). DEGs (red) are FDR<0.05 and (log2FC)> 0.2. Exemplary “consolidation-dependent” DEGs are labeled in black. b) Number of DEGs per neuron subtype. Heatmap of the number of shared DEGs between each neuron subtype. c) Heatmap of the log2 fold change of exemplary “consolidation-dependent” DEGs (FR vs NF) per neuron subtype. Each gene is further labeled with annotations of interest. d) The percentile in which a TRAPed neuron in C0 lies in the distribution of expression of a C0 DEG (log2cpm space). Box plots show the the log2cpm distribution for each C0 DEG. Hierarchical clustering shows one common transcriptional program that is concertedly upregulated. e) Fraction of cells in each neuron subtype that are activated with the transcriptional program (i.e. DEGs) from C0.

To begin to understand the biological significance of these genes, we applied a set of strict criteria that were selected to identify the most probable candidate effectors. First, each gene had to be differentially expressed in at least ¾ of biological replicates, enforcing reproducibility. Subsequent removal of DEGs that are also differentially expressed between the inactive populations in FR vs NF mice allowed the identification of changes specific to active populations (**Extended Fig 4a**). Next, DEGs must be differentially expressed when FR cells are compared to NR and HC controls, ensuring that DEGs are not just a consequence of a fear experience (**Extended Fig 4b**). Lastly, DEGs further passed a permutation test with shuffled labels. These criteria produced a set of 102 “consolidation-specific DEGs” which were biologically robust and remote memory consolidation-specific (see *Methods)* (**Fig 3c**). Several genes encoded for proteins with regulatory roles, including known transcriptional regulators *Slc30a9, Hmg20a, Hnrnpk, Zfp706*, as well as translation regulating factors *Nck2, Alpl1, and Eif2ak1*. Interestingly, even among the condensed list of consolidation-specific DEGs, we find strong enrichments in processes including the regulation of: vesicle exocytosis (*Vamp2, Gdi2, Rab15, Rab5a, Rab24, Atp6v0c, Syt13, Stx1b, Nsf*), trans-membrane transport (*Slc25a46, Mfsd14a, Tmem50a, Gpm6a, Mfsd14b, Abcf3*), dendritic spine density (*Strip1, Pls3, Gsk3b*), and long-range intracellular protein transport (*Timm29, Atad1, Pak1, Plehkb2, Sarnp, Rtn3, Dmtn, Sar1a, Hid1*) (**Fig 3c, Extended Fig 3b-c**). While these processes are highly linked to synaptic plasticity, we have identified distinct genes out of an expansive plasticity-related coding space that may dictate the specificity in which two neurons communicate during the maintenance of a remote fear-memory trace. Lastly, more than half of consolidation-specific DEG candidates have known associations with neuronal diseases in the Malacards database, suggesting links between the functional role of these genes to various neuronal dysfunctions (Alzheimer Disease, Neuroblastoma, Schizophrenia) and the regulation of long-term memory.

Single-cell information also allowed us to probe the diversity of transcriptional programs within each subtype. When cells were clustered by the percentile of their expression level for each DEG within the range of all TRAPed cells, distinct populations of “highly activated” and “lowly activated” cells emerged. This suggests that one transcriptional program is concertedly upregulated and perhaps even co-regulated, during memory consolidation (**Fig 3d**). To determine the subtype-specificity of transcriptional programs, individual neurons from each subtype were assigned an “activation state” (see *Methods*). A cell is considered to be “activated” if 50% of the subtype-specific DEGs tested is expressed above the level of the 90th percentile of the distribution in NF controls. Indeed, the fraction of cells activated with the subtype-specific DEGs was generally highest in the corresponding subtype when compared to the activation levels in other subtypes or in the inactive populations (**Fig 3e, Extended Fig 5a**). Together, this could indicate the presence of subtype-specific common regulatory elements.

To address this possibility, we analyzed our DEGs using Hypergeometric Optimization of Motif Enrichment (HOMER) to search for common regulatory motifs in an unbiased manner. The search was performed anywhere from −400 to +100 bp of the transcription start-site for each DEG (**Extended Fig 5b**). We found 12 putative *de novo* and 2 known motifs enriched within our target gene set (p>0.01). Unexpectedly, dependencies on *CREB, NFKb, CBP, C/EBP*, AP-1 - canonical transcriptional regulators of neuronal activity, plasticity and short-term memory retrieval (<24h post-learning)^27–29^ - were absent. In particular, the *HIF1b* binding motif was found in >40% of our target DEG sequences, including synaptic transmission and plasticity-related genes *Rab5a, Rab24, Vamp2, Gdi2, Gpm6a, Strip1, Ptp4a1, Trim32, Mfsd14a, Mfsd14b, and Slc25a46*. Interestingly, these findings are in line with recent work supporting a potential dual role for *HIF-1* transcription factors during hippocampal-dependent spatial learning and early consolidation under normoxic conditions^30^.

## Consolidation involves a sustained increase in specific presynaptic vesicle fusion and exocytosis genes

To further elucidate the significance of these consolidation-dependent transcriptional programs, we used STRING to look for known and predicted protein-protein interactions. *K-means* clustering of the gene nodes revealed a significantly connected network (p=1.75e-06) that was centered around a large cluster of genes encoding for proteins for vesicle-mediated transport, exocytosis and neurotransmitter secretion, all of which were highly connected (confidence=0.4, **Extended Fig 5c**). Remarkably, 20/102 consolidation-specific DEGs fall within these functional categories. These DEGs include syntaxin-1b (*Stx1b*), synaptotagmin-13 (*Syt13*), vesicle-associated membrane protein 2 (*Vamp2*), vesicle-fusing ATPase (*Nsf*), and ras-related protein (*Rab5a*), all of which individually are functionally linked to the SNARE-complex and to vesicle exocytosis at the presynaptic terminal (**Fig 4a**). Interestingly, the two most highly and ubiquitously upregulated genes across subtypes were *Serinc-1* and *Serinc-3*, recently discovered to be serine incorporators^31^. Upon delivering serine to the endoplasmic reticulum, phosphatidylcholine and serine are used to synthesize phosphatidylserine (PS), a component of the inner synaptic membrane. Notably, PS phospholipids were first characterized as a binding partner for synaptotagmins upon exposure to Ca2+^32–35^, which suggests that *Serinc-1* and -*3* may have important roles in enhancing PS levels and facilitating vesicle membrane fusion during memory consolidation. Finally, *in situ* hybridization confirmed the endogenous proportions of neuronal subtypes in TRAPed populations (**Extended Fig 6a-b**), as well as the upregulated expression of key consolidation-specific genes including *Serinc3, Syt13, Vamp2,* and *Stx1b* within respective neuron subtypes (**Fig 4b-c, Extended Fig 6c**).

**Figure 4.**
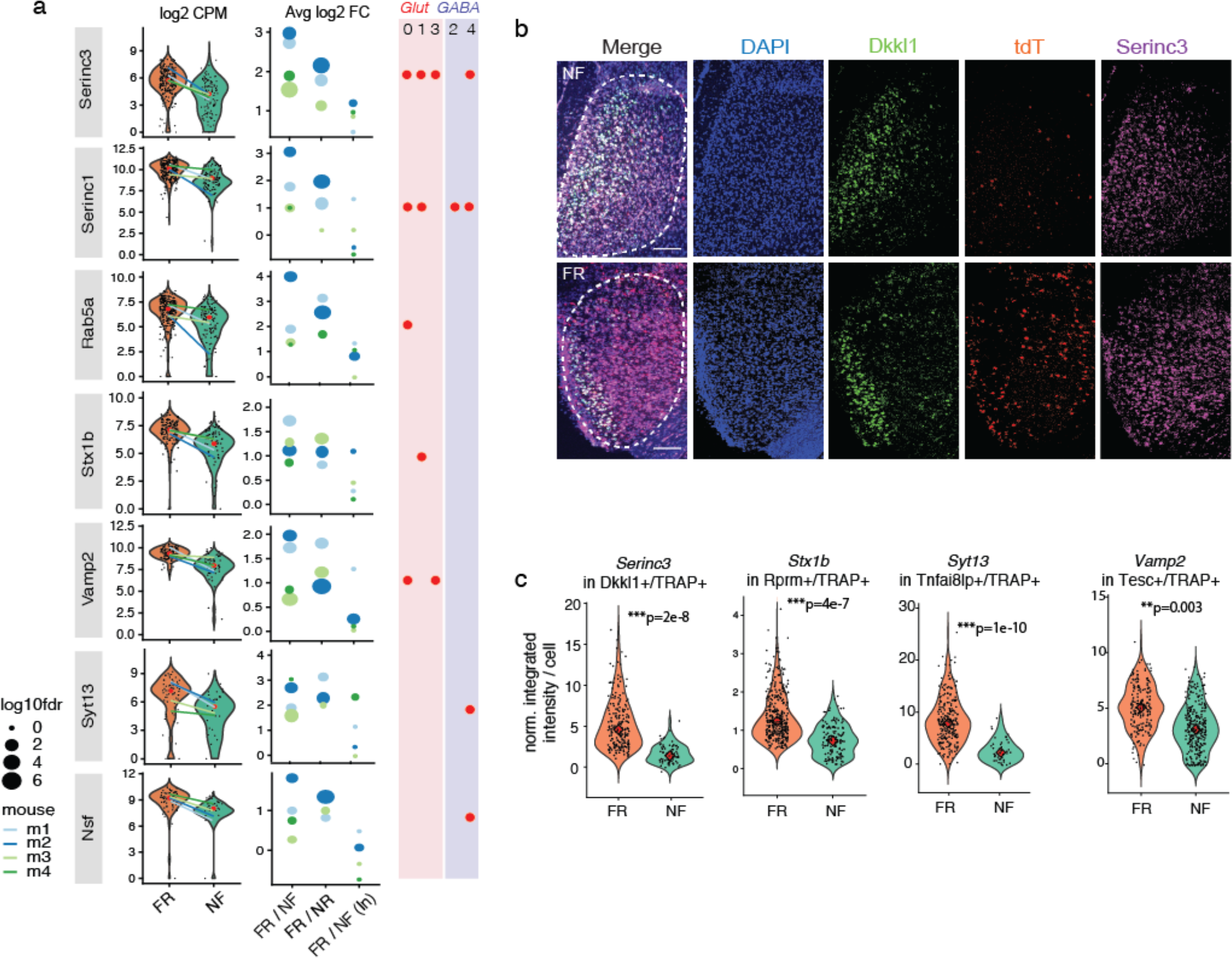
Memory consolidation involves the upregulation of genes coding for specific vesicle exocytosis proteins. a) Violin plots of consolidation-dependent DEGs (from Fig 3c) that are known to regulate vesicle membrane fusion and exocytosis at the pre-synaptic terminal. Violin plots are overlayed with a drumstick plot indicating the average expression per mouse. Bubble plots depict the log2 fold change and degree of significance per mouse. In addition to FR/NF, DEGs were also confirmed to be differentially expressed when compared to the NR control (FR/NR), and specific only to the active populations, as indicated by the comparison between inactive populations (FR/NF (In)). Red dots indicate which neuronal subtype these particular vesicle exocytosis genes are upregulated. b) Representative *in situ* images (RNAscope) of *Serinc3* in *Dkkl1*+/*tdT*+ cells in FR and NF mice. Dotted lines outline the mPFC region. Scale bars = 100 micron. c) *In situ* validation of key vesicle exocytosis genes in various neuron subtypes, including *Serinc3* (in *Dkkl1*+ subtype, n=228 (FR) and 92 (NF) cells over 3 mice/condition), *Stx1b* (in *Rprm*+ subtype, n=342 (FR) and 143 (NF) cells over 3 mice/condition), *Syt13* (in *Tnfai8lp*+ subtype, n=288 (FR) and 52 (NF) cells over 3 mice/condition), and Vamp2 (in *Tesc*+ subtype, n=356 (FR) and 325 (NF) cells over 3 mice/condition).

While synaptic plasticity is known to be required for consolidation, the molecular and cellular mechanisms through which plasticity is achieved can vary widely depending on the experience, brain region and neuron type. Although such diversity is expected, technological limitations in accessing memory engrams and the methods to characterize the transcriptomic landscape of single neurons in a high-throughput manner has limited understanding this diversity. Here, we show for the first time that *(1)* enhanced membrane-fusion and vesicle-exocytosis may be a critical mode of synaptic strengthening during memory consolidation, *(2)* a specific-set of exocytosis-related genes is found to be involved out of a vast coding-space which could allow highly unique, experience-specific connections to be made, and *(3)* these particular transcriptional programs are sustained and thus likely required to maintain the memory trace weeks after learning.

## Non-neuronal cells exhibit significant transcriptional changes that are distinct from neurons

Remarkably, we discovered that non-neuronal cells also exhibited sustained transcriptional changes upon memory consolidation (FR compared to NF mice, **Fig 5a-b, Extended Fig 7a-b**). These signatures were distinct from neurons, indicating that distinct non-neuronal programs may exert supporting roles in the maintenance of the remote fear-memory trace. Surprisingly, in all cell types more than 95% of these DEGs were upregulated, suggesting an overall transcriptional activation during consolidation. Not only is this response sustained weeks after the initial learning, it is detected even without enrichment of the non-neuronal cells directly associated with the TRAPed engram cells since the TRAP method is neuron-specific. Thus, it is likely that a robust response of non-neuronal cells contributes to memory consolidation.

**Figure 5.**
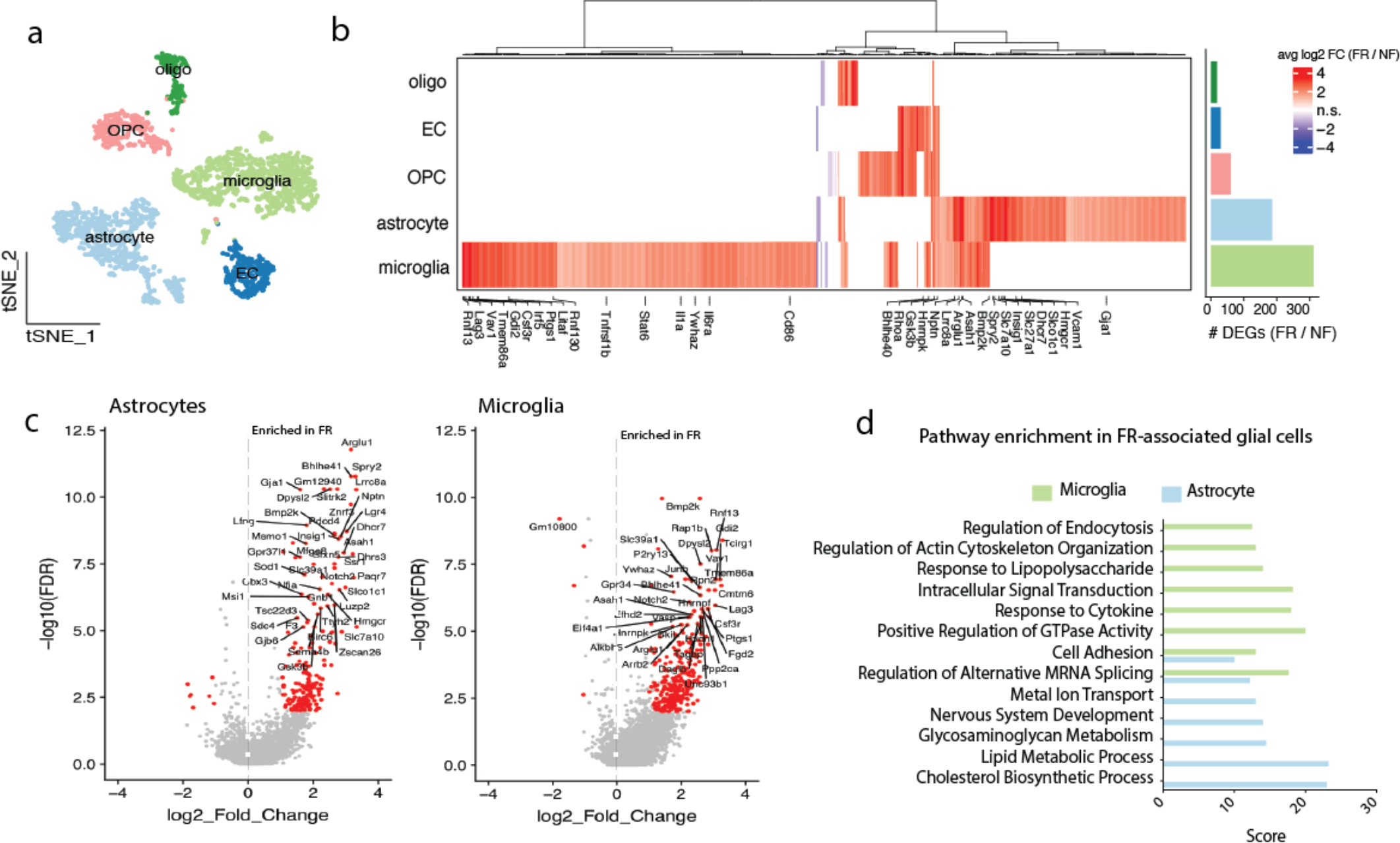
Non-neuronal cells exhibit significant transcriptional changes that are distinct from neurons. a) tSNE of all non-neuronal cells collected (collected in an unbiased manner from tdT− gates) reveal 5 cell types (astrocyte, endothelial, microglia, OPCs and oligodendrocytes). b) Heatmap of DEGs in non-neuronal cells (FR over NF). DEGs are defined as log2FC>1 and FDR<0.01. Fold change of genes that do not pass this threshold are not displayed (white = not significant). Genes are clustered via the ward.D algorithm. Bar graph depicts the number of DEGs that satisfy this criteria in each non-neuronal cell type. Top DEGs for glial cells (astrocytes and microglia) that are also differentially expressed in FR vs NR, are labeled. c) Volcano plots of DEGs up and downregulated in astrocytes and microglia in FR vs NF mice. DEGs (FDR>0.01, log2FC>1) are labeled in red, and exemplary DEGs (high log2FC and −log10FDR) are labeled in black. d) GO biological function pathway analysis of the DEGs in microglia and astrocytes

We find that the upregulated transcriptional programs are unique to each cell type. Of 605 total unique DEGs, only 100 are shared between two cell types or more. Astrocytes and microglia showed the greatest intersection and number of transcriptional changes, with 181 and 308 genes perturbed, respectively, in FR mice (log2FC>1 and FDR<0.01) (**Fig 5c**). Most of these DEGs represent largely diverging pathways (**Fig 5d**). In particular, upregulated astrocytic genes were enriched in lipid, cholesterol and steroid metabolic functions (*Gja1*, *Hmgcr, Dhcr7, Insig1, Acsl3, Idi1, Acsbg1, 10Asah1, Hacd3*) as well as glucose transport (*Abcc5, Slc39a1, Slc6a1, Slc27a1, Slco1c1, Gnb1, Ttyh1*), suggesting that enhanced metabolic support from astrocytes could be required during the neuronal consolidation process. This is particularly interesting given that astrocytes have long been known to support the immensely high energy requirements of neurons, and that astrocyte-neuron metabolic coupling is elevated during neuronal activity^36,37^. Moreover, 95/181 astrocyte DEGs were also reproduced when comparing FR to NR mice, suggesting that a large portion of these effects is specific to the recall-experience itself and not merely a remnant of the fear experience.

In contrast, DEGs from microglial cells were enriched in innate immunity (*Il6r, Stat6, Csf3r, Il1a, Irf5, Cd86, Tnfrsf1b, Ywhaz, Litaf, Ptgs1, Gdi2, Rnf13*) and cytoskeletal re-organization / focal adhesion maintenance (*Cdc42, Rhoa, Rhoh, Prkcd, Vasp, Arf6, Vav1, Actr2*) pathways, suggesting that upregulation of specific inflammatory molecules and enhancement of cell migration in microglia may be involved in supporting the maintenance of the memory trace. While less is known regarding the immunomodulatory roles of microglia in memory and learning, previous work has shown that low levels of inflammatory cytokines (such as IL-1, IL-6 and TNF-alpha) can regulate neuronal circuit remodeling and long-term potentiation^38,39^.

In addition to neuron-neuron coupling, additional communication programs between neurons and non-neuronal cells may be acquired to support the memory trace over long periods. We looked for the expression of receptors or ligands in non-neuronal cells whose known binding partner^40^ is perturbed in TRAPed FR neurons (**Extended Fig 8a-b**). In particular, we focused on genes that were differentially expressed in both the ligand-bound and receptor-bound cell type with fear-memory consolidation (**Extended Fig 8c**). Interestingly, in FR mice, we find the upregulation of neuronal neurogilins-1 and -3 (*Ngln1, Ngln3*) and its binding partner neurexin-1 (*Nrxn1*) on astrocytes. A family of CAMs that is a critical component of the bipartite synapse, neuroligin-neurexin complexes are known to act to enhance neuron-glia adhesions and modulate synaptic function^41–43^. Thus, the concerted upregulation of these binding pairs in FR mice strongly suggests a role of astrocyte-neurexin-neurogilin interactions in the maintenance of synaptic strength during fear memory storage. Altogether, we show for the first time that non-neuronal cells exhibit consolidation-dependent transcriptional perturbations and that microglia and astrocytes may play particularly important roles via a distinct set of genes that support the remote memory trace.

## Discussion

While high-resolution gene-expression atlases of the brain have provided invaluable information regarding cellular taxonomy ^44–46^, characterization of activity- and experience-dependent cell states within these cell types (including non-neuronal cells) is necessary for understanding how experience modulates synaptic plasticity and neuronal circuitry. In particular, the molecular mechanisms underlying the storage of information within neuronal ensembles, and how precisely gene transcription operates in the conversion of short- to long-term memories, is largely unclear. Here, using a combination of activity-dependent labeling of neurons (TRAP2), contextual fear-conditioning, and single cell transcriptomics, we discover the unexpected existence of sustained and coordinated transcriptional programs within activated neuronal ensembles that likely contribute to long-term memory storage in the mPFC.

Activity-driven transcription is well established and known to shape cortical function and plasticity^47^. A wide range of stimuli including acute sensory experience, metabolic changes, stress, injury and pharmacological intervention evoke new gene expression programs, typically through the action of activity-dependent IEGs (e.g. *c-Fos, Arc, Jun, Egr1, Egr2, Homer1)* and TFs that regulate them (including *CREB, SRF, CRE, Nf-kB, C/EBP*). Induction of these early-response genes (ERGs) can directly alter synaptic transmission ^47–50^. Recently, scRNA-seq of neurons in the visual cortex 1h after light-stimulus confirmed the presence of canonical stimulus-dependent ERGs such as *Nr4a1, Nr4a2, Fos, Fosl2, Egr1, Junb*, but also found considerable divergences within the ERG programs across different neuronal and non-neuronal cell types^51^. Moreover, transcriptional profiles of neurons 4 h post-stimulation were significantly different, resulting in diverse late response gene (LRG) programs that included secreted neuropeptides and synaptic organizers (e.g. *Cbln4, Crh*). Intriguingly, the gene expression programs we found upregulated during memory consolidation (~3 weeks post-learning) are largely non-intersecting with these well-established IEGs that are upregulated immediately following salient novel behavioral experience. While canonical IEGs and ERTFs were likely active during the initial fear-learning, they are nearly absent 25 days post-shock (9 days after retrieval). Instead, the sustained transcriptional changes we discovered are likely experience- and region-specific LRG-like programs that are acquired gradually to consolidate a long-term memory trace. Together, this suggests that the activity-dependent transcriptional landscape cannot merely be generalized across experiences, brain regions, nor time.

In contrast to transcriptional programs immediately induced by acute sensory experience, those involved in more gradual processes like memory storage are much less understood. A cascade of molecular changes is thought to strengthen ‘engram’ synapses in multiple brain regions, including the amygdala, mPFC, and hippocampus. These changes likely depend on gene-expression, as evidenced by the sensitivity of memory consolidation to broad transcriptional inhibitors^52,53^. Thus, identifying such gene-expression changes provides a gateway to understanding memory storage. A microarray study in the dentate gyrus revealed the dynamic nature of short-term consolidation (0-24 h post- passive avoidance learning), which is characterized by a perturbation of ~500 genes involved in translation/transcription initiation, cell adhesion, neurotransmission and intercellular trafficking^54^. Recently, Rao-Ruiz et al^55^ isolated engram-specific hippocampal neurons for single-cell sequencing 24 h after contextual fear conditioning and found an enrichment of activity-regulated genes (e.g. *Arc, Npas4, Dusp1, Atf3, Cdkn1a*), many of which were CREB-dependent. However, these studies were restricted to recent memory (retrieval at <24 h after learning), leaving the mechanisms of remote memory unexplored. In fact, new neural pathways are known to be recruited in cortical structures such as the mPFC over time, suggesting a time-dependent evolution of gene programs as one transitions to remote memory. Indeed, we find that the majority of early activity-dependent gene programs perturbed in activated neurons during recent fear memory retrieval^55^, associative fear-learning^56^, post-visual stimulus,^51^ or novel environment exposure^57^ are not found within our memory consolidation DEGs (**Extended Figure 9a-c**). In fact, none of plasticity-related vesicle exocytosis genes we identified were found in these data sets, suggesting the uniqueness of these signature to remote memory consolidation.

Interestingly, TRAPed neurons in mice with fear but no recall (NR) injected with 4-OHT 16d after the fear-conditioning also exhibited sustained transcriptional changes at moderate levels (when compared to NF) (**Extended Fig 10**). However, these DEGs are largely non-intersecting with consolidation-specific DEGs, suggesting that (1) the process of recall further induces new transcriptional programs in a different set of neurons and (2) the experience of fear itself can induce long-lasting gene expression programs.

It is well established that learning and short-term memory (<24 h post-learning) involves CREB-dependent gene networks ^27,28^. A recent study found that a subset of mPFC neurons activated (and labeled) during fear-learning is involved in remote memory expression and CREB-dependent^58^. However, detailed analysis of memory consolidation DEGs and their upstream regulators did not reveal any canonical activity-regulated transcription factors such as CREB, CBP, NF-kB, AP-1, or C/EBP^10,29,59–61^. This suggests that the subset of mPFC neurons activated during remote retrieval may differ from those activated during learning, and that they operate under other regulators. While this does not preclude the requirement of CREB in long term memory storage, our data is the first to show that remote memory storage mechanisms could be governed by a different set of transcriptional and regulatory programs than in learning or recent-memory consolidation (<24 h).

The ability to form and maintain unique synaptic connections that encode a particular memory out of a vast pool of other experiences requires a highly complex coding-space that is likely both molecular and physical in nature. Indeed, the ability of all mPFC neuron types to be activated during consolidation (**Fig 2f)**and the heterogeneity of the activated transcriptional programs (**Fig 3c**) suggests that discrete neuronal populations play differential roles in maintaining the memory trace and thus expand the coding-space. In particular, genes involved in membrane fusion and vesicle exocytosis are strongly upregulated and distributed across subtypes, including *Vamp2, Rab5a, Nsf, Stx1b, Syt13, Rab15, Gdi2, Rtn3, and Sar1a*, a number of which are known members of the SNARE-complex^62,63^. While none of these genes have yet to be described in relation to remote-memory, their increased expression here could indicate particular roles in regulating neurotransmitter release throughout memory consolidation. We also uncover a potentially novel role of *Serinc-1* and -*3* in memory storage, which could function to enhance membrane fusion via Ca2+-dependent synaptotagmins. Interestingly, phosphatidylserine (PS) supplements have been long been purported to aid in aging-related memory loss, with the basis that aging is correlated to loss of PS in the mammalian brain^64,65^. The heterogeneity of transcriptional programs between neuron types and the coordination of specific plasticity-related genes points towards the diversity of plasticity mechanisms during memory consolidation. Taken together, our findings provide important insight into the transcriptional basis of memory consolidation and shed light on the therapeutic potential of targeting consolidation-dependent gene-expression programs to address memory loss or enhancement in neuronal disorders.

## Acknowledgements

We thank Drs. Lu Chen, Mu Zhou, and Jie Li for discussion of the experimental design; S. Kolluru and D. Henderson for assistance in library preparation; N. Neff and J. Okamoto for assistance with sequencing; J. Lui for advice on brain dissociation; Drs. L. Denardo, J. Lui and L. Luo for the gifting and help with TRAP2 line; W. Wang, G. Stanley, F. Horns for helpful discussions and computational assistance. S.R.Q. is a Chan Zuckerberg Investigator.

## Author Contributions

XJ and TCS designed animal experiments. MBC and SRQ designed scRNA-seq experiments. XJ performed animal experiments and brain dissection. MBC performed brain dissociation, flow cytometry, single-cell library preparation and sequencing pipelines. XJ performed in *situ* hybridization and imaging. MBC performed all scRNA-seq data and image analysis, with input from XJ, SRQ and TCS. MBC wrote the manuscript with significant contributions from XJ, SRQ, and TCS. TCS and SRQ oversaw the project.

## METHODS

### Mice

All animal experiments were conducted following protocols approved by the Administrative Panel on Laboratory Animal Care at Stanford University. TRAP2; Ai14 mouse line was kindly gifted from Luo lab at Stanford^21,23^. TRAP2 mice were heterozygous for the Fos2A-iCreER allele, and homozygous for Ai14. Mice were group-housed (maximum 5 mice per cage) on a 12 h light–dark cycle (7 am to 7 pm, light) with food and water freely available. Male mice 42–49 days of age were used for all the experiments. Mice were handled daily for 3 days before their first behavior experiment.

### Genotyping

The following primers: (For) GAG GGA CTA CCT CCT GTA CC, (Rev) TGC CCA GAG TCA TCC TTG GC were used for genotyping of the Fos2A-iCreER allele.

### Fear conditioning

The fear conditioning training was performed as previously described^66^. Briefly, mice were individually placed in the fear conditioning chamber (Coulbourn Instruments) located in the center of a sound attenuating cubicle, which was cleaned with 10% ethanol to provide a background odor. A ventilation fan provided a background noise at ~55 dB. After a 2 min exploration period, three tone-footshock pairings separated by 1 min intervals were provided. The 85 dB 2 kHz tone lasted for 30 s, and the footshocks were 0.75 mA and lasted for 2 s. The foot shocks co-terminated with the tone. The mice remained in the training chamber for another 60 s before being returned to the home cages. For the context recall, mice were placed back into the original conditioning chamber for 5 min 16 days after the training. The behavior of the mice was recorded with the FreezeFrame software and analyzed with the FreezeView software (Coulbourn Instruments). Motionless bouts lasting more than 1 s were considered as freeze. Data were analyzed with the tracking software Viewer III (Biobserve).

### TdTomato florescence examination

Mice were deep anesthetized with tribromoethanol and perfused with PBS followed by fixative (4% paraformaldehyde diluted in PBS). The brains were then removed and postfixed in 4 °C overnight and immersed in 30% sucrose solution for 2 days before being sectioned at 50 μm thicknesses on a cryostat (CM3050 S; Leica Biosystems). Imaging was performed with a scanning microscope (BX61VS; OLYMPUS CORPORATION).

### Single-cell dissociation and flow cytometry

mPFC regions were micro-dissected from live vibratome sections (300 um thick) of the prefrontal cortex. Tissue pieces were enzymatically dissociated via a papain-based digestion system (Worthington, Cat # LK003150). Briefly, tissue chunks were incubated in 1mL of papain (containing L-cysteine and EDTA), DNAse, and kyneurenic acid for 1 hour at 37C and 5% CO2. After 10 min of incubation, tissues were triturated briefly with a P1000 wide bore pipette tip and returned. Cells were triturated another 4 times (~30 each) with a P200 pipette tip over the rest of the remaining incubation time. At room temperature, cell suspensions were centrifuged at 350g for 10 min, resuspended in 1mL of EBSS with 10% v/v ovomucoid inhibitor, 4.5% v/v DNAse and 0.1% v/v kyneurenic acid, and centrifuged again. Supernatant was removed and cells 1mL ACSF was added. ACSF was composed of: 1mM KCl, 7mM Mgcl2, 0.5 mM Cacl2, 1.3 mM NaH2PO4, 110 mM choline chloride, 24mM NaHCO3, 1.3 mM Na Ascorbate, 20mM glucose and 0.6mM sodium pyruvate. Cells were passed through a 70 um cell strainer to remove debris. Hoescht stain was added (1:2000, Life Technologies, Cat #H3570) and incubated in the dark at 4C for 10 min. Samples were centrifuged (350g for 8 min at 4C) and resuspended in 0.5mL of ACSF and kept on ice for flow cytometry.

Cells were sorted via the Sony SH800 into 96 or 384 well plates (Biorad) directly into lysis buffer^67^ with oligodT, and immediately snap frozen until processing. A positive “TRAP” gate was set for cells that were both Hoescht+ and tdT+. A negative “TRAP” gate was set for all Hoecht+ and tdT− cells in general. No gating on forward or backscatter was used to avoid size biases that might be present in a heterogenous neuronal population. Each plate was kept on the sorter for <25 min to prevent evaporation.

### Sequencing

Cell lysis, first-strand synthesis and cDNA synthesis was performed using the Smart-seq-2 protocol as described previously^67^ in both 96-well and 384-well formats, with some modifications. After cDNA amplification (23 cycles), cDNA concentrations were determined via capillary electrophoresis (96-well format) or the PicoGreen quantitation assay (384-well format) and wells were cherry-picked to improve quality and cost of sequencing. Cell selection was done through custom scripts and simultaneously normalizes cDNA concentrations to ~0.2 ng/uL per sample, using the TPPLabtech Mosquito HTS and Mantis (Formulatrix) robotic platforms. Libraries were prepared, pooled and cleaned using the Illumina Nextera XT kits or in-house Tn5, following the manufacturer’s instructions. Libraries were then sequenced on the Nextseq or Novaseq (Illumina) using 2 × 75bp paired-end reads and 2 × 8bp index reads with a 200 cycle kit (Illumina, 20012861). Samples were sequenced at an average of 1.5M reads per cell.

### RNA scope

RNAscope experiment was performed following the manufactory’s instructions using RNAscope multiplex fluorescent reagent kit v2 (ACD Cat #323100). All probes were purchased from existing stocks of custom designed from ACD.

### Bioinformatics and data analysis

#### Mapping to the genome

Sequences from the Nextseq or Novaseq were demultiplexed using bcl2fastq, and reads were aligned to the mm10 genome augmented with ERCC sequences, using STAR version 2.5.2b. Gene counts were made using HTSEQ version 0.6.1p1. All packages were called and run through a custom Snakemake pipeline. We applied standard algorithms for cell filtration, feature selection, and dimensionality reduction. Briefly, genes appearing in fewer than 5 cells, samples with fewer than 100 genes, and samples with less than 50,000 reads were excluded from the analysis. Out of these cells, those with more than 30% of reads as ERCC, and more than 10% mitochondrial or 10% ribosomal were also excluded from analysis. Counts were log-normalized then scaled where appropriate.

Next, the *Canonical Correlation Analysis* function from the Seurat package ^68^ was used to align raw data from multiple experiments. Only the first 10 Canonical Components (CCs) were used. After alignment, relevant features were selected by filtering expressed genes to a set of ~2500 with the highest positive and negative pairwise correlations. Genes were then projected into principal component space using the robust principal component analysis (rPCA). Single cell PC scores and genes loadings for the first 20 PCs were used as inputs into Seurat’s (v2) *FindClusters* and *RunTsne* functions to calculate 2-dimensional tSNE coordinates and search for distinct cell populations. Briefly, a shared-nearest-neighbor graph was constructed based on the Euclidean distance metric in PC space, and cells were clustered using the Louvain method. Cells and clusters were then visualized using 3-D t- distributed Stochastic Neighbor embedding on the same distance metric. A neuron is characterized as “TRAPed” trapped if it satisfies 2 conditions: (1) from the tdT+ sort gate (tdT protein positive) (2) tdT mRNA raw count > 0. Differential gene expression analysis was done by applying the Mann-Whitney U-test on various cell population. Raw p-values were adjusted via the false discovery rate (FDR). Permutation tests were then performed on all genes of interest. All graphs and analyses were generated and performed in R. GeneAnalytics and GeneCards- packages offered by Gene Set Enrichment Analysis (GSEA) tool was used for GO/KEGG/REACTOME pathway analysis and classification of enriched genes in each subpopulation.

#### Finding “consolidation-specific DEGs”

To reduce our list of DEGs (FR TRAP vs NF TRAP results in 1291 DEGs, cells from 4 biological replicates pooled, logFC>0.3, FDR<0.05) to only the most recall-specific, 4 steps were taken. First, DEGs are re-calculated by assessing each experiment individually using the whole transcriptome, and only DEGs that intersect in ¾ replicates are kept. While this decreases the statistical power, it ensures biological reproductivity. The 3 out of 4 criteria was chosen as a compromise due to the high strictness of 4 out of 4 which yielded only 7 DEGs. All resulting DEGs are found in the initial DEG list (all replicates pooled). Second, “inactive” (tdT negative) populations were also compared (FR inactive vs NF inactive) and any DEGs which were intersecting with the DEGs left after the first criteria, were removed. This ensures that DEGs are activity-dependent, and not merely an overall upregulation in all cells due to the experience. This routinely removed genes such as *Hsp90aa1* and *Pcna-ps2*. Third, the remaining DEGs had to be differentially expressed when FR TRAP was compared to either FR TRAP and HC TRAP. This ensures that the DEGs are specific to only neuronal ensembles that labeled by memory recall, and not due to forms of baseline activity (HC) or activity remaining from the initial fear learning (NR). Last, the remaining DEGs must pass a permutation test where the training labels are shuffled and a distribution of log2FC is computed based on these labels. The true observed logFC must be above the 95^th^ percentile of the distribution of the shuffled distribution. After placing these constraints, 102 genes remain from the original list of 1291.

#### Assessment of “Activation Score”

A TRAPed (or inactive) cell is considered to be “activated” by the consolidation-DEG program if 25, 50 or 75% of the subtype-specific DEGs (“consolidation-specific DEGs” only) is expressed above the 90th percentile of the distribution of that gene in NF TRAP controls from the same subtype. This calculation is then repeated with DEG programs specific to each neuronal subtype. The fraction of cells activated with the subtype-specific signature is calculated as the number of “activated” cells divided by all cells in the subtype/activity group.

#### Regulatory Motif analysis

Known and de novo motifs enrichment was found using HOMER by inputting the list of 102 consolidation-specific DEGs and using the function *findMotifs.pl* and the criteria ‘–*start −400 -end 100 -len 8,10 -p 2*’. Motif locations in specific DEGs were found using the *-find* <*motif file*> option of *findMotifs.pl*.

#### RNAscope image analysis

Images were taken at using a Nikon Confocal Microscope (at 10X or 20X, NA=0.45) and images were processed in ImageJ to only obtain the mPFC regions. The resulting images were fed into a custom image analysis pipeline was developed on CellProfiler (using a combination of the functions *IdentifyPrimaryObjects, RelateObjects, FilterObjects, MeasureObjectIntensity, ClassifyObjects,* and *CalculateMath*. Custom pipeline in SI Methods). Briefly, images were corrected with control slides (unstained sample and negative control probes) and cells were segmented using the DAPI signal. Those harboring a signal (above a set threshold level) for both the subtype marker and tdT probe were retained. The integrated fluorescence intensity of the DEG probe was calculated for each DAPI+/Subtype+/tdT+ cell. Cells that were not double-positive were not considered. The integrated fluorescent intensity was then normalized to the integrated DAPI signal per cell and results were plotted with custom scripts in R.

## References

1. Lechner, H. A., Squire, L. R., Byrne, J. H. & Mu, G. Remembering Müller and Pilzecker. 77–88

2. Müller, G. E. & Pilzecker, A. Experimentelle beiträge zur lehre vom gedächtniss. (J.A. Barth, 1900).

3. Squire, L. R. Mechanisms of memory. Science (80-.). 232, 1612 LP – 1619 (1986).

4. Hebb, D. Organization of Behavior: A Neuropsychological Theory. John Wiley Sons Inc, NJ (1941). doi:10.1177/000271625027100159

5. Agranoff, B. W., Davis, R. E., Casola, L. & Lim, R. Actinomycin D Blocks Formation of Memory of Shock-Avoidance in Goldfish. Science (80-.). 158, 1600 LP – 1601 (1967).

6. Flexner, J. B., Flexner, L. B. & Stellar, E. Memory in Mice as Affected by Intracerebral Puromycin. Science (80-.). 141, 57 LP – 59 (1963).

7. Taubenfeld, S. M., Milekic, M. H., Monti, B. & Alberini, C. M. The consolidation of new but not reactivated memory requires hippocampal C/EBPβ. Nat. Neurosci. 4, 813–818 (2001).

8. Kandel, E. R. The Molecular Biology of Memory Storage : A Dialogue Between Genes and Synapses. 294, 1030–1039 (2015).

9. Flexner, L. B. & Flexner, J. B. Effect of acetoxycycloheximide and of an acetoxycycloheximide-puromycin mixture on cerebral protein synthesis and memory in mice. 55, 369–374 (1966).

10. Alberini, C. M. Transcription factors in long-term memory and synaptic plasticity. Physiol. Rev. 89, 121–145 (2009).

11. Alberini, C. M. & Kandel, E. R. The Regulation of Transcription in Memory Consolidation. Cold Spring Harb. Perspect. Biol. 7, (2015).

12. Donahue, C. P. et al. Transcriptional profiling reveals regulated genes in the hippocampus during memory formation. Hippocampus. 12, 821–833 (2002).

13. Tonegawa, S., Morrissey, M. D. & Kitamura, T. The role of engram cells in the systems consolidation of memory. Nat. Rev. Neurosci. 19, 485–498 (2018).

14. Reijmers, L. G., Perkins, B. L., Matsuo, N. & Mayford, M. Localization of a Stable Neural Correlate of Associative Memory. Science (80-.). 317, 1230 LP – 1233 (2007).

15. Tonegawa, S., Liu, X., Ramirez, S. & Redondo, R. Memory Engram Cells Have Come of Age. Neuron. 87, 918–931 (2015).

16. DeNardo, L. & Luo, L. Genetic strategies to access activated neurons. Curr. Opin. Neurobiol. 45, 121–129 (2017).

17. Allen, W. E. et al. Thirst-associated preoptic neurons encode an aversive motivational drive. Science (80-.). 357, 1149 LP – 1155 (2017).

18. Guenthner, C. J., Miyamichi, K., Yang, H. H., Heller, H. C. & Luo, L. Permanent genetic access to transiently active neurons via TRAP: targeted recombination in active populations. Neuron. 78, 773–784 (2013).

19. DeNardo, L. A. et al. Temporal evolution of cortical ensembles promoting remote memory retrieval. Nat. Neurosci. 22, 460–469 (2019).

20. Semon, R. W. Die Mneme als erhaltendes Prinzip im Wechsel des organischen Geschehens. . (Engelmann, 1911).

21. Thompson, R. F. In Search of Memory Traces. Annu. Rev. Psychol. 56, 1–23 (2004).

22. Josselyn, S. A., Köhler, S. & Frankland, P. W. Finding the engram. Nat. Rev. Neurosci. 16, 521 (2015).

23. Minatohara, K., Akiyoshi, M. & Okuno, H. Role of Immediate-Early Genes in Synaptic Plasticity and Neuronal Ensembles Underlying the Memory Trace. Front. Mol. Neurosci. 8, 78 (2016).

24. Euston, D. R., Gruber, A. J. & McNaughton, B. L. The role of medial prefrontal cortex in memory and decision making. Neuron. 76, 1057–1070 (2012).

25. Bontempi, B., Laurent-Demir, C., Destrade, C. & Jaffard, R. Time-dependent reorganization of brain circuitry underlying long-term memory storage. Nature 400, 671–675 (1999).

26. Zeisel, A. et al. Molecular Architecture of the Mouse Nervous System. Cell 174, 999–1014.e22 (2018).

27. Suzuki, A. et al. Upregulation of CREB-Mediated Transcription Enhances Both Short- and Long-Term Memory. J. Neurosci. 31, 8786 LP – 8802 (2011).

28. Kida, S. et al. CREB required for the stability of new and reactivated fear memories. Nat. Neurosci. 5, 348–355 (2002).

29. Gass, P. et al. Deficits in memory tasks of mice with CREB mutations depend on gene dosage. Learn. Mem. 5, 274–288 (1998).

30. C O’Sullivan, N., Sheridan, G. & Murphy, K. Transcriptional Profiling of Hippocampal Memory-Associated Synaptic Plasticity: Old Friends and New Faces. in Transcription Factors CREB and NF-kB: Involvement in Synaptic Plasticity and Memory Formation 43–65 (Bentham Science Publishers, 2012). doi:10.2174/978160805257811201010043

31. Inuzuka, M., Hayakawa, M. & Ingi, T. Serinc, an Activity-regulated Protein Family, Incorporates Serine into Membrane Lipid Synthesis. J. Biol. Chem. 280, 35776–35783 (2005).

32. Perin, M. S., Friedt, V. A., Mignery, G. A., Jahn, R. & Ii, T. C. S. Phospholipid binding by a synaptic vesicle protein homologous to the regulatory region of protein kinase C. 345, 260–263 (1990).

33. Brose, N., Petrenko, A. G., Sudhof, T. C. & Jahntt, R. Synaptotagmin : A Calcium Sensor on the Synaptic Vesicle Surface. 256, 1021–1026 (1992).

34. Zhang, X., Rizo, J. & Südhof, T. C. Mechanism of Phospholipid Binding by the C2A-Domain of Synaptotagmin I. Biochemistry. 37, 12395–12403 (1998).

35. Davis, A. F. et al. Kinetics of Synaptotagmin Responses to Ca^2+^ and Assembly with the Core SNARE Complex onto Membranes. Neuron. 24, 363–376 (1999).

36. Pellerin, L. & Magistretti, P. J. Glutamate uptake into astrocytes stimulates aerobic glycolysis: a mechanism coupling neuronal activity to glucose utilization. Proc. Natl. Acad. Sci. U. S. A. 91, 10625–10629 (1994).

37. Bélanger, M., Allaman, I. & Magistretti, P. J. Brain Energy Metabolism: Focus on Astrocyte-Neuron Metabolic Cooperation. Cell Metab. 14, 724–738 (2011).

38. Williamson, L. L., Sholar, P. W., Mistry, R. S., Smith, S. H. & Bilbo, S. D. Microglia and Memory: Modulation by Early-Life Infection. J. Neurosci. 31, 15511 LP – 15521 (2011).

39. Yirmiya, R. & Goshen, I. Immune modulation of learning, memory, neural plasticity and neurogenesis. Brain. Behav. Immun. 25, 181–213 (2011).

40. Ramilowski, J. A. et al. A draft network of ligand–receptor-mediated multicellular signalling in human. Nat. Commun. 6, 7866 (2015).

41. Knight, D., Xie, W. & Boulianne, G. L. Neurexins and Neuroligins: Recent Insights from Invertebrates. Mol. Neurobiol. 44, 426–440 (2011).

42. Hillen, A. E. J., Burbach, J. P. H. & Hol, E. M. Cell adhesion and matricellular support by astrocytes of the tripartite synapse. Prog. Neurobiol. 165–167, 66–86 (2018).

43. Südhof, T. C. Synaptic Neurexin Complexes: A Molecular Code for the Logic of Neural Circuits. Cell. 171, 745–769 (2017).

44. Zeisel, A. et al. Cell types in the mouse cortex and hippocampus revealed by single-cell RNA-seq. Science (80-.). 347, 1138 LP – 1142 (2015).

45. Saunders, A. et al. Molecular Diversity and Specializations among the Cells of the Adult Mouse Brain. Cell 174, 1015–1030.e16 (2018).

46. Rosenberg, A. B. et al. Single-cell profiling of the developing mouse brain and spinal cord with split-pool barcoding. Science (80-.). 360, 176 LP – 182 (2018).

47. West, A. E. & Greenberg, M. E. Neuronal Activity–Regulated Gene Transcription in Synapse Development and Cognitive Function. Cold Spring Harb. Perspect. Biol. (2011).

48. Shepherd, J. D. & Bear, M. F. New views of Arc, a master regulator of synaptic plasticity. Nat. Neurosci. 14, 279 (2011).

49. Sheng, M. & Greenberg, M. E. The regulation and function of c-fos and other immediate early genes in the nervous system. Neuron. 4, 477–485 (1990).

50. Chang, M. C. et al. Narp regulates homeostatic scaling of excitatory synapses on parvalbumin-expressing interneurons. Nat. Neurosci. 13, 1090 (2010).

51. Hrvatin, S. et al. Single-cell analysis of experience-dependent transcriptomic states in the mouse visual cortex. Nat. Neurosci. 21, 120–129 (2018).

52. Chen, D. Y. et al. A critical role for IGF-II in memory consolidation and enhancement. Nature 469, 491–U63 (2011).

53. Igaz, L. M., Vianna, M. R. M., Medina, J. H. & Izquierdo, I. Two Time Periods of Hippocampal mRNA Synthesis Are Required for Memory Consolidation of Fear-Motivated Learning. J. Neurosci. 22, 6781 LP – 6789 (2002).

54. O’Sullivan, N. C. et al. Temporal change in gene expression in the rat dentate gyrus following passive avoidance learning. J. Neurochem. 101, 1085–1098 (2007).

55. Rao-Ruiz, P. et al. Engram-specific transcriptome profiling of contextual memory consolidation. Nat. Commun. 10, 1–14 (2019).

56. Cho, J.-H., Huang, B. S. & Gray, J. M. RNA sequencing from neural ensembles activated during fear conditioning in the mouse temporal association cortex. Sci. Rep. 6, 31753 (2016).

57. Lacar, B. et al. Nuclear RNA-seq of single neurons reveals molecular signatures of activation. Nat. Commun. 7, 11022 (2016).

58. Matos, M. R. et al. Memory strength gates the involvement of a CREB-dependent cortical fear engram in remote memory. Nat. Commun. 10, 1–11 (2019).

59. Ahn, H. J. et al. c-Rel, an NF-κB family transcription factor, is required for hippocampal long-term synaptic plasticity and memory formation. Learn. Mem. 15, 539–549 (2008).

60. Bourtchuladze, R. et al. Deficient long-term memory in mice with a targeted mutation of the cAMP-responsive element-binding protein. Cell. 79, 59–68 (1994).

61. Sakamoto, K., Karelina, K. & Obrietan, K. CREB: a multifaceted regulator of neuronal plasticity and protection. J. Neurochem. 116, 1–9 (2011).

62. Ungermann, C. & Langosch, D. Functions of SNAREs in intracellular membrane fusion and lipid bilayer mixing. J. Cell Sci. 118, 3819 LP – 3828 (2005).

63. Südhof, T. C. & Rothman, J. E. Membrane Fusion: Grappling with SNARE and SM Proteins. Science (80-.). 323, 474 LP – 477 (2009).

64. Kato-Kataoka, A. et al. Soybean-derived phosphatidylserine improves memory function of the elderly Japanese subjects with memory complaints. J. Clin. Biochem. Nutr. 47, 246–255 (2010).

65. Glade, M. J. & Smith, K. Phosphatidylserine and the human brain. Nutrition. 31, 781–786 (2015).

66. Zhou, M. et al. A central amygdala to zona incerta projection is required for acquisition and remote recall of conditioned fear memory. Nat. Neurosci. 21, 1515–1519 (2018).

67. Picelli, S. et al. Full-length RNA-seq from single cells using Smart-seq2. Nat. Protoc. 9, 171–181 (2014).

68. Butler, A., Hoffman, P., Smibert, P., Papalexi, E. & Satija, R. Analysis Integrating single-cell transcriptomic data across different conditions, technologies, and species. Nat. Biotechnol. 36, 411–420 (2018).

