## Extended Figures for "Persistent transcriptional programs are associated with remote memory in diverse cells of the medial prefrontal cortex"

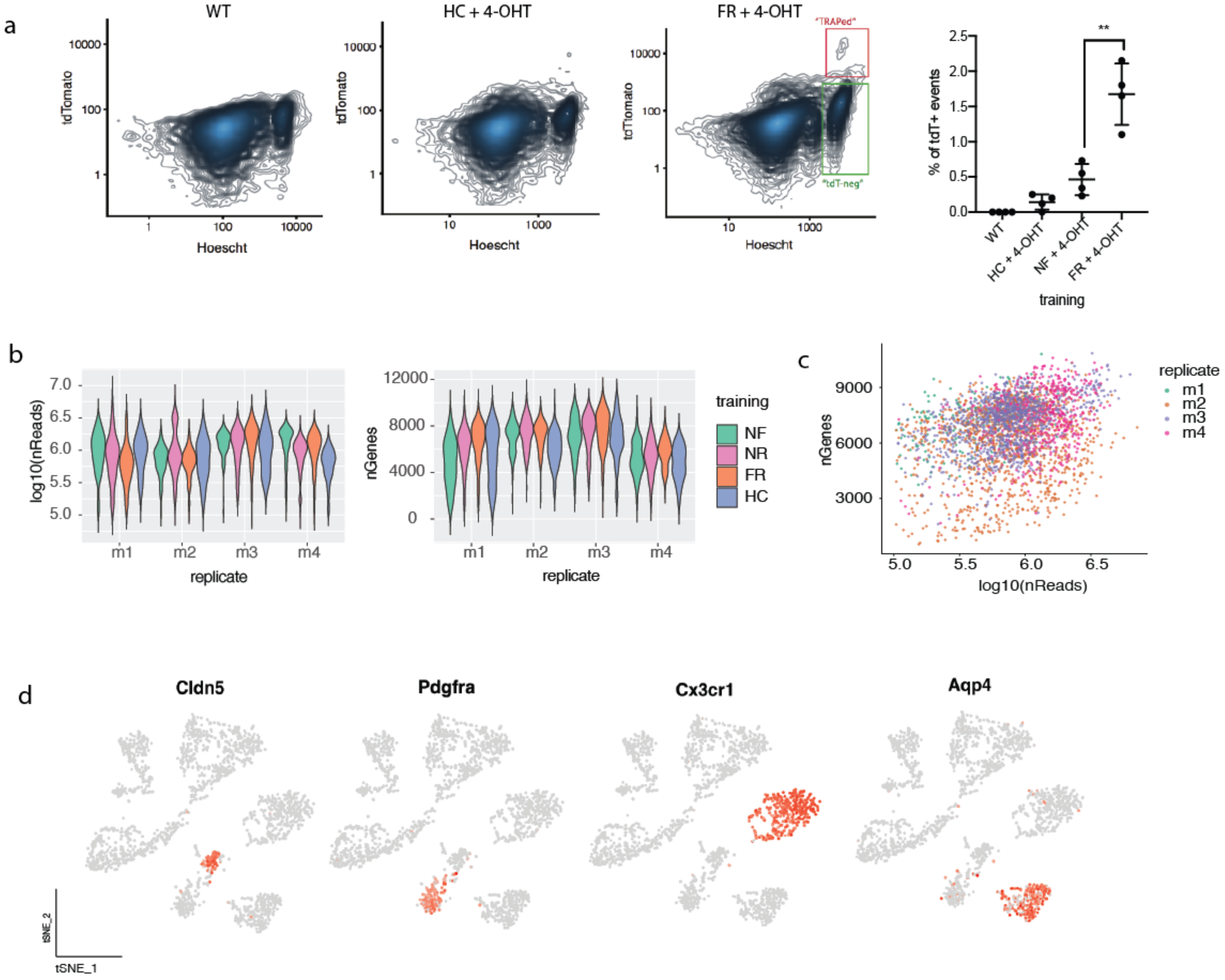


**Extended Fig 1 (related to Fig 1). Flow cytometry, sequencing quality and identification of major cell types**

a) Representative FACs plot of the amount of tdT+ events per training condition. In scatter plot, each point represents one mouse.
b) Violin plots of the number of reads and number of genes per biological replicate (m1-m4) for each training paradigm, in neurons.
c) Scatterplot of the number of genes detected and number of reads obtained per cell for all training conditions combined.
d) Scaled expression of canonical markers in non-neuronal cells (*Cldn5*-BECs, *Pdgfra*-OPCs, *Cx3cr1*-microglia, *Aqp4*-astrocytes)

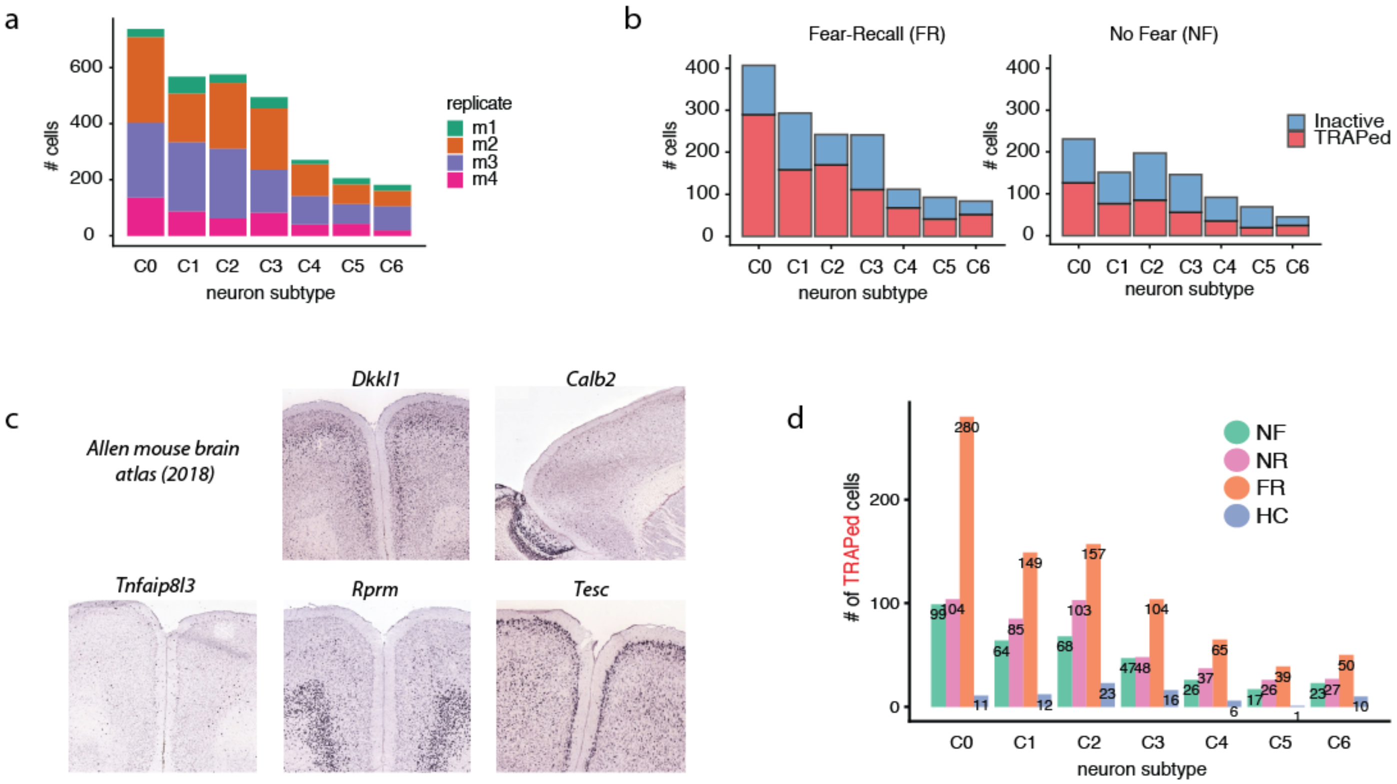


**Extended Fig 2 (related to Fig 2). Cell numbers per neuronal subtype**

a) Number of cells from each biological replicate that were annotated as one of 7 defined neuron subtypes (C0-C6)

b) Number of TRAPed and Inactive cells (as defined by non-zero expression of *tdT* mRNA) collected per neuron subtype, in either fear-recall (FR) or no-fear (NF) mice.

c) Representative images of subtype marker genes in the Allen Brain ISH atlas.

d) Number of TRAPed (tdT mRNA+) cells collected in each experimental condition that fall in one of 7 neuronal subtype categories.

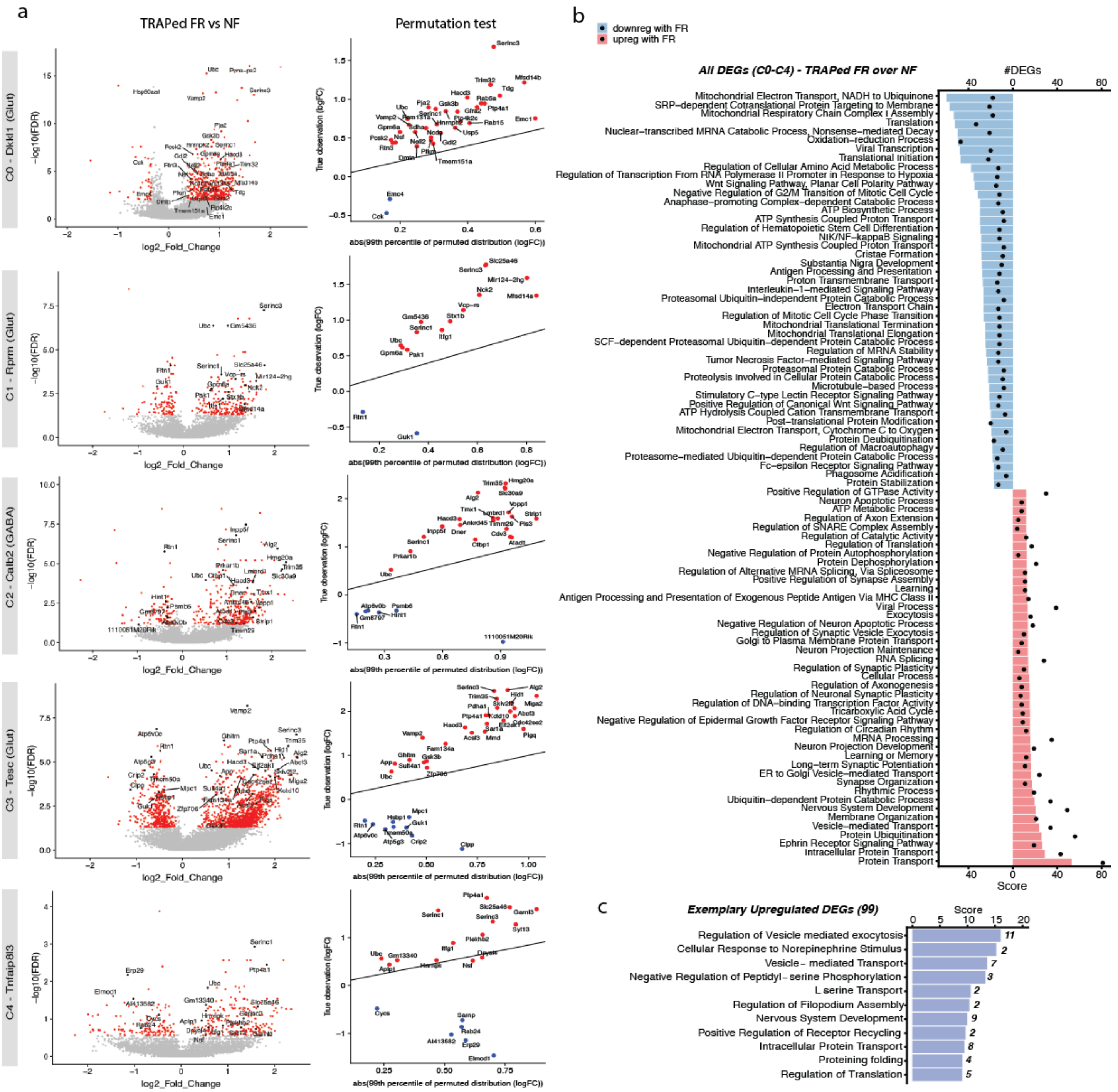


**Extended Fig 3 (related to Fig 3). Differential gene expression in neuronal subtypes (FR vs NF TRAP)**

a) (Left) Volcano plots of DEGs in FR vs NF mice for each neuron subtype. DEGs found when all replicates are pooled are shown in red. Exemplary consolidation-dependent DEGs are labeled in black. (Right) Permutations are performed for every consolidation-dependent DEG for each neuronal population. Upregulated DEGs lying above the y=x line (red) and downregulated DEGs lying below the y=x line (blue) are considered to be above the 99$^{th}$ percentile of the permuted distribution.

b) GO enrichment analysis of all up- and down-regulated DEGs (941 DEGs up, 384 DEGs down, all neuron subtypes combined) when all replicates are pooled. Bars show the enrichment scores (GeneAnalytics) for the GO pathway and dots indicates the number of DEGs involved.

c) GO enrichment analysis of only the upregulated consolidation-dependent DEGs from all neuron subtypes. Number indicates number of consolidation-dependent DEGs involved in that pathway.

a

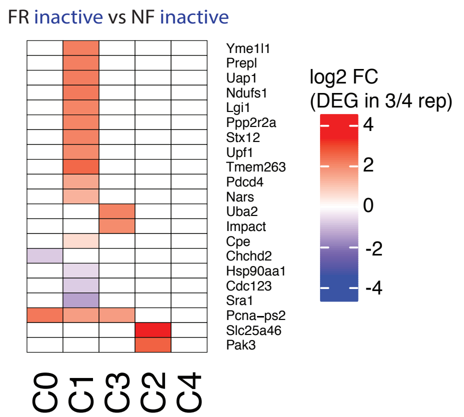


b

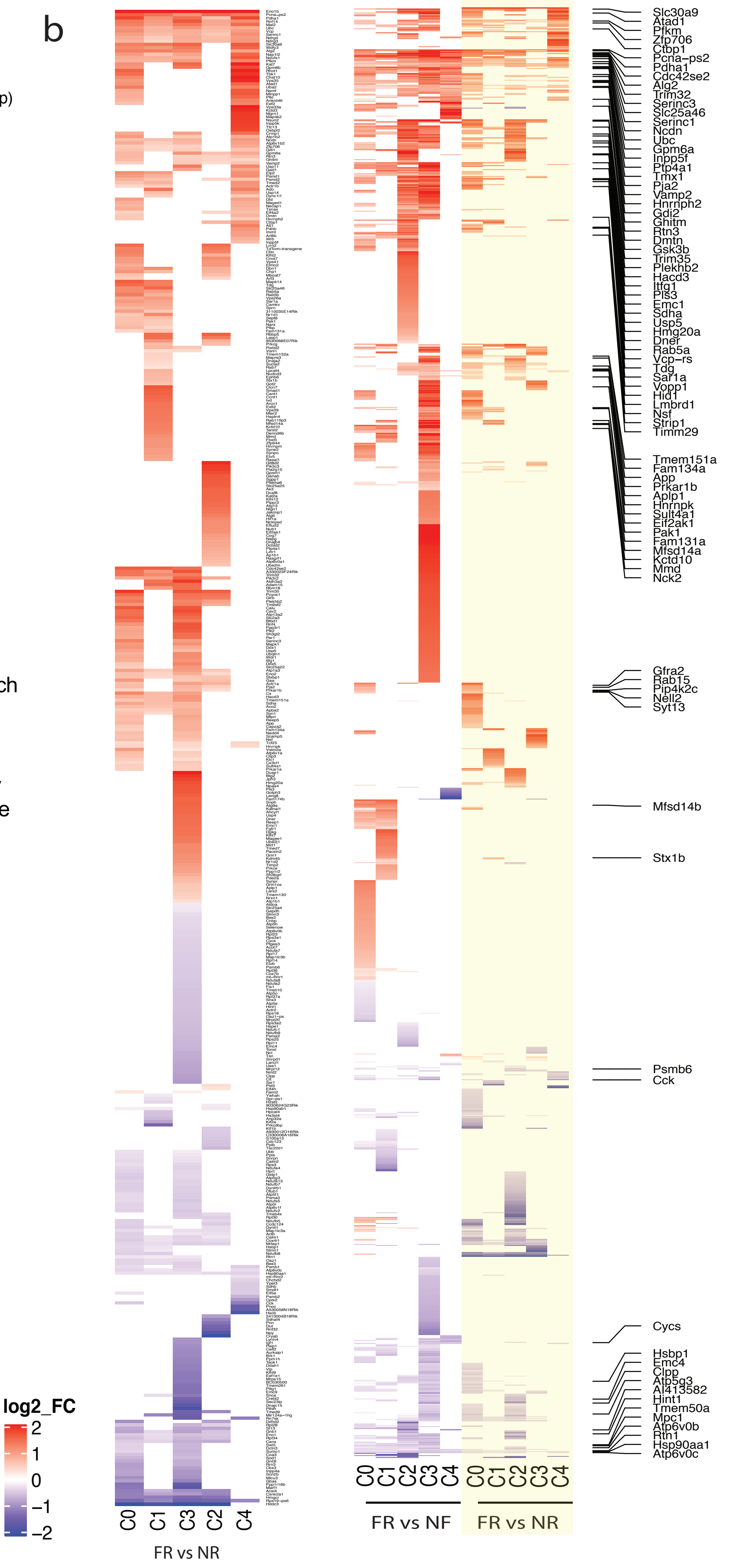


**Extended Fig 4 (related to Fig 3). DEGs resulting from comparisons with other training conditions**

a) DEGs in each neuron subtype when inactive (tdT-) neurons are compared between FR and NF mice.

b) (Left) log2 fold change of DEGs for each subtype when FR is compared to NR mice (no recall). (Right) The union of DEGs for FR vs NF, and FR vs NR (yellow). Genes labeled are exemplary consolidation-dependent DEGs that are differentially expression in both comparisons.

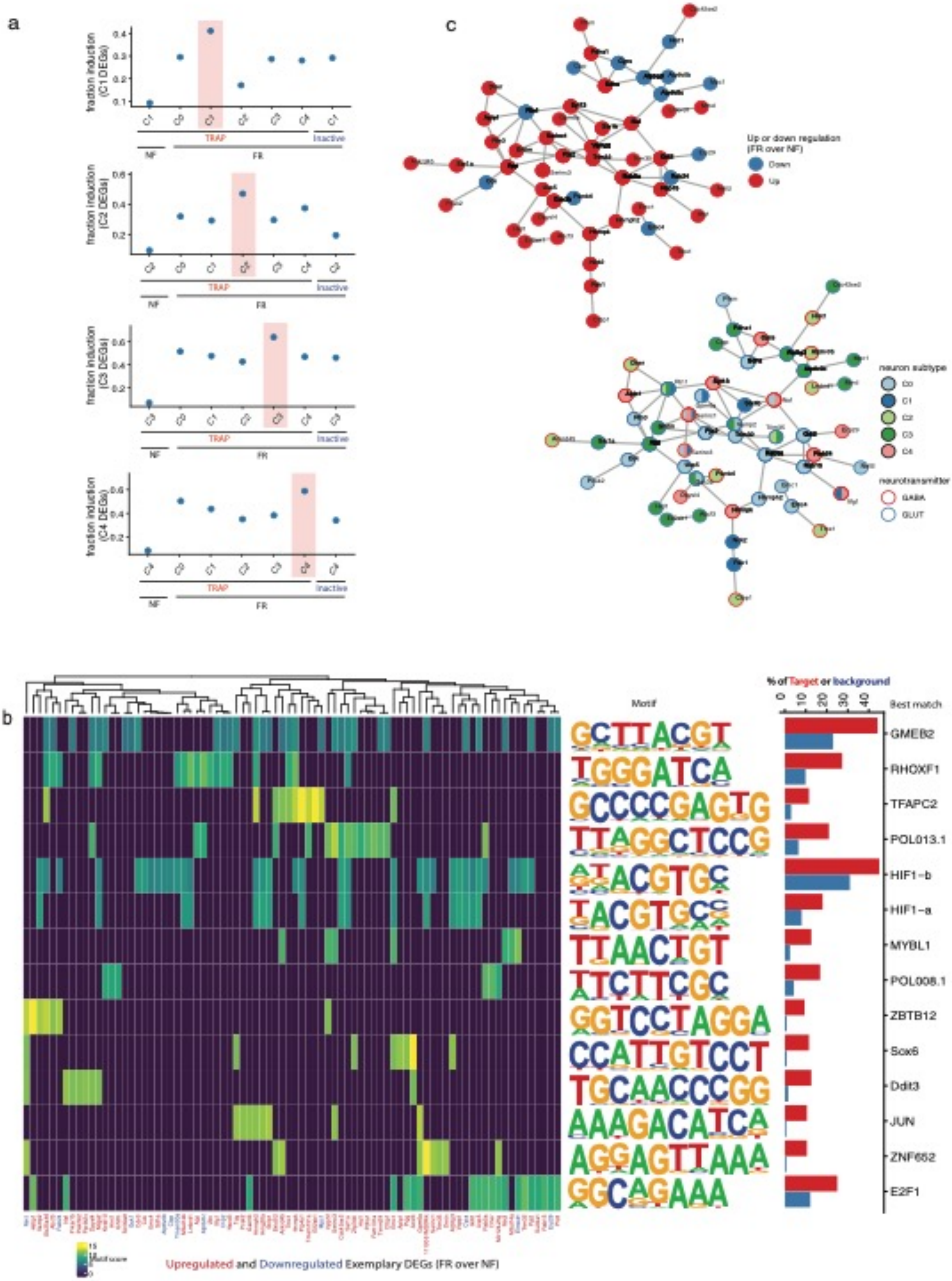
a
fraction induction (C1 DEGs)
fraction induction (C2 DEGs)
fraction induction (C3 DEGs)
fraction induction (C4 DEGs)
TRAP
Inactive
NF
FR
b
Motif
% of Target or background
Best match
GMEB2
RHOXF1
TFAPC2
POL013.1
HIF1−b
HIF1−a
MYBL1
POL008.1
ZBTB12
Sox6
Ddit3
JUN
ZNF652
E2F1
Upregulated and Downregulated Exemplary DEGs (FR over NF)
c
Up or down regulation (FR over NF)
Down
Up
neuron subtype
C0
C1
C2
C3
C4
neurotransmitter
GABA
GLUT

**Extended Fig 5 (related to Fig 3) – Analysis of common transcription programs, regulatory genes and networks**

a) Fraction of cells in each neuron subtype that are induced with the transcriptional program (i.e. DEGs) from a neuron subtype. Overall, the activation program of each TRAPed neuron subtype is found to be more specific to it than the inactive population, or other neuron subtypes.

b) (Left) De novo regulator motif discovery: analysis was performed using HOMER on the subset of 102 exemplary consolidation-dependent DEGs (neuronal, FR / NF) by looking at the sequences -400 to +100 bp from the TSS. 12 *de novo* and 2 known motifs were found (only motifs with an enrichment p-value < 1e-2) were kept) . Heatmap depicts the "motif score" of each DEG for each motif , and genes and motifs were clustered via the ward.D method. (Right) Bar graph depicting the % of the DEGs (target sequences) that possess a match for the motif within -400 to +100 bp from the TSS, vs the % of background sequences. For de novo motifs, the best match gene is listed on the right. HIF1b and HIF1a are matches to known motifs.

c) (Top) Hypothesize protein-protein interactions of a subset of consolidation-dependent DEGs (TRAPed FR/NF) using the STRING database (https://string-db.org/). Only genes that are connected at a confidence level of 0.4 (medium) are shown. Connections indicate a possible existence of an interaction between two proteins. Genes are colored by up of down-regulation in FR/NF. (Bottom) Same network plot, with nodes colored by the neuron subtype which differentially regulates the DEG.

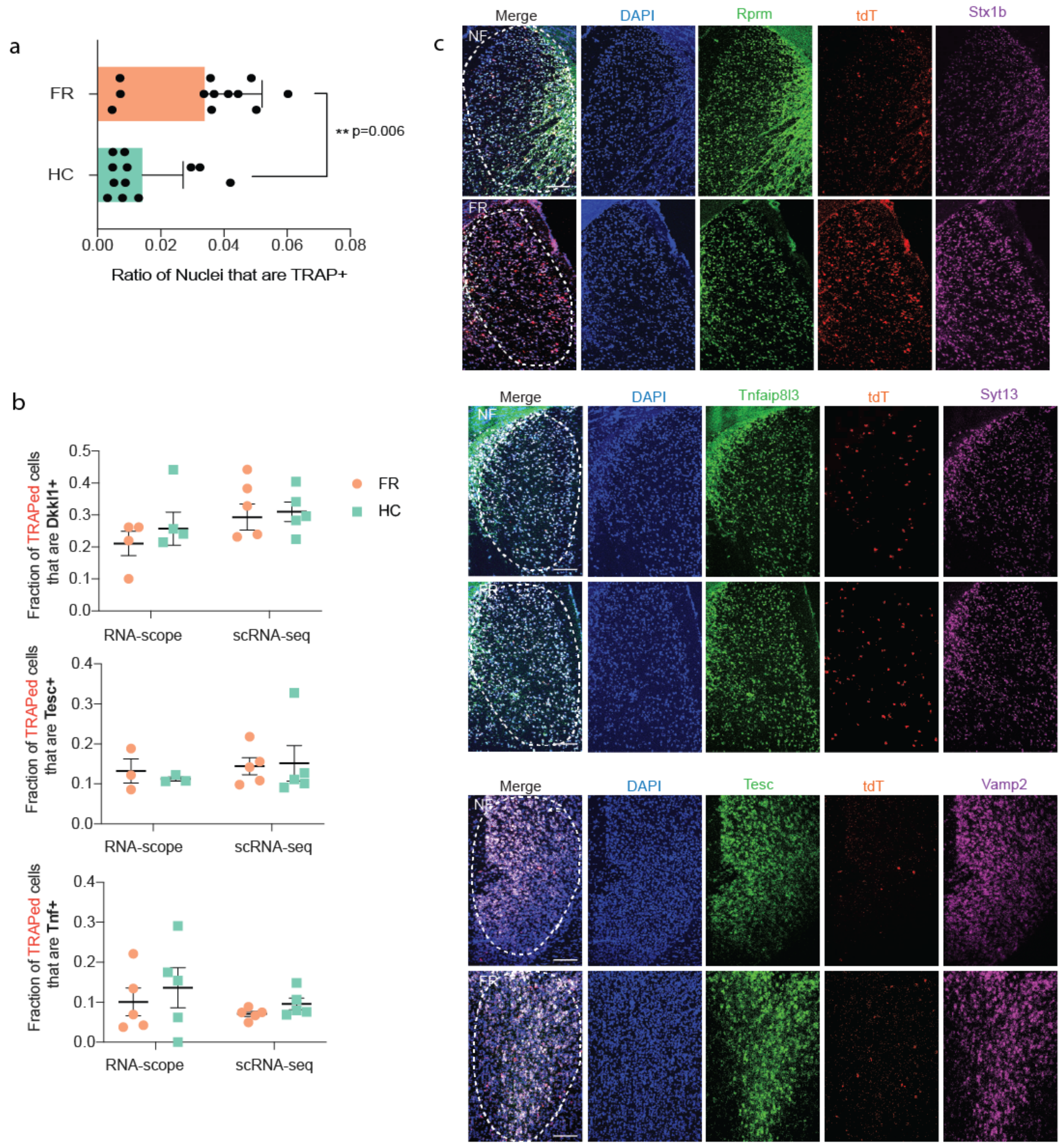


**Extended Fig 6 (related to Fig 4). In situ validation of tdT levels, neuronal subtypes and consolidation-dependent DEGs**

a) Ratio of nuclei (cells) that are tdT+ (as detected by in situ staining) per training condition. Each data point represents one region of interest (spread over 3 mice/condition).

b) Ratio of TRAPed cells that are positive for a neuronal subtype marker obtained either via the RNA-scope method, or by scRNAseq. TRAPed cells are defined as DAPI+/tdT+ in RNAscope quantification, and as tdT mRNA count >1 in scRNA-seq (post-QC). No significant differences are found between FR and HC within either RNA-scope or scRNA-methods.

c) RNAscope *in situ* validation of key consolidation-dependent DEGs (Stx1b in Rprm+/tdT+ cells, Syt13 in Tnfaip8l3+/tdT+ cells, Vamp2 in Tesc+/tdT+ cells)

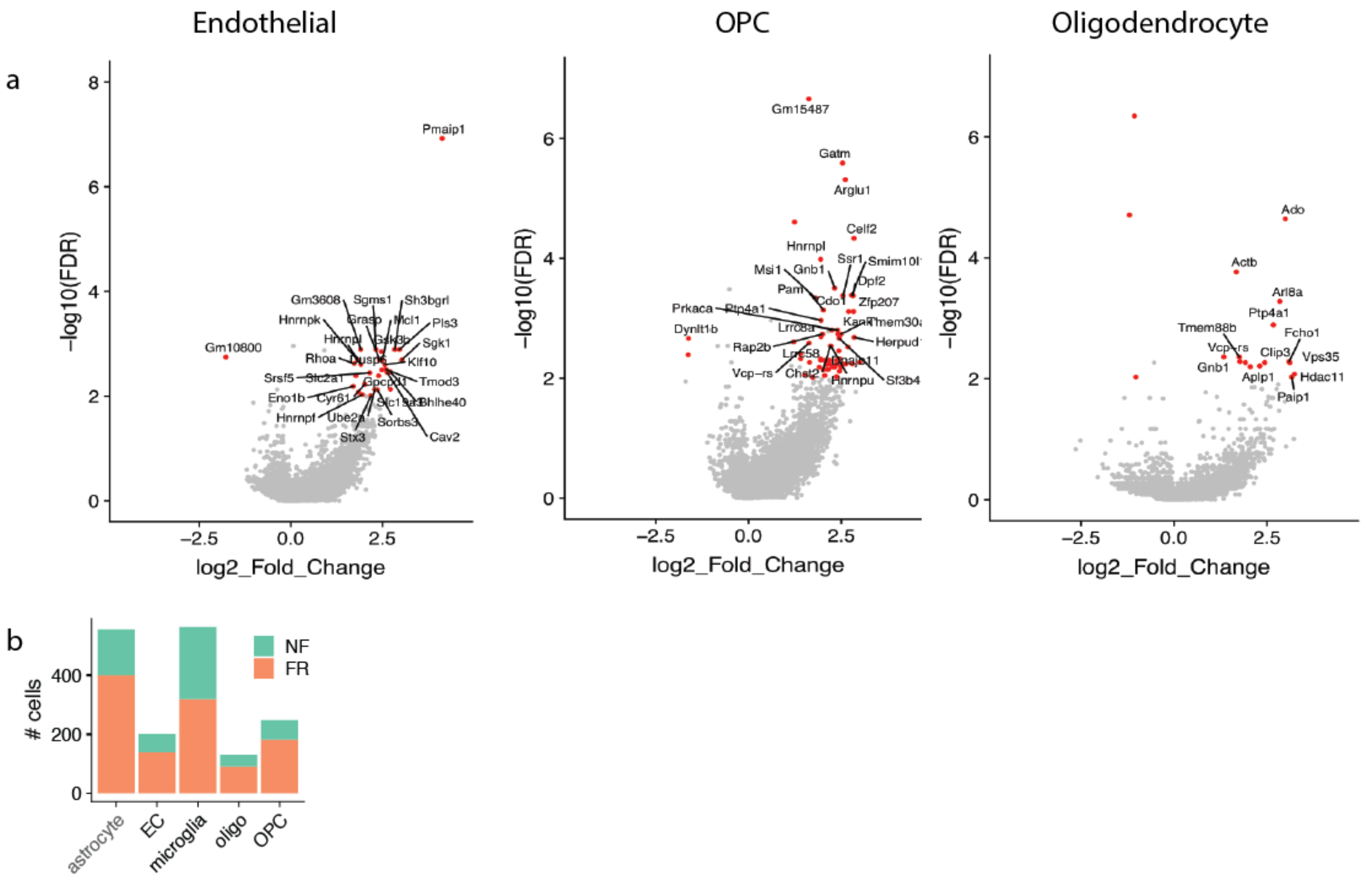


**Extended Fig 7 (related to Fig 5). Differentially expressed genes in non-neuronal cells**

a) Volcano plots of non-neuronal cell types when comparing cells in FR vs NF mice. DEGs (FDR>0.01, log2FC>1) are labeled in red, and exemplary DEGs (high log2FC and log10FDR) are labeled in black.

b) Bar graph of the number of non-neuronal cells collected in this study, for each cell type and experimental condition.

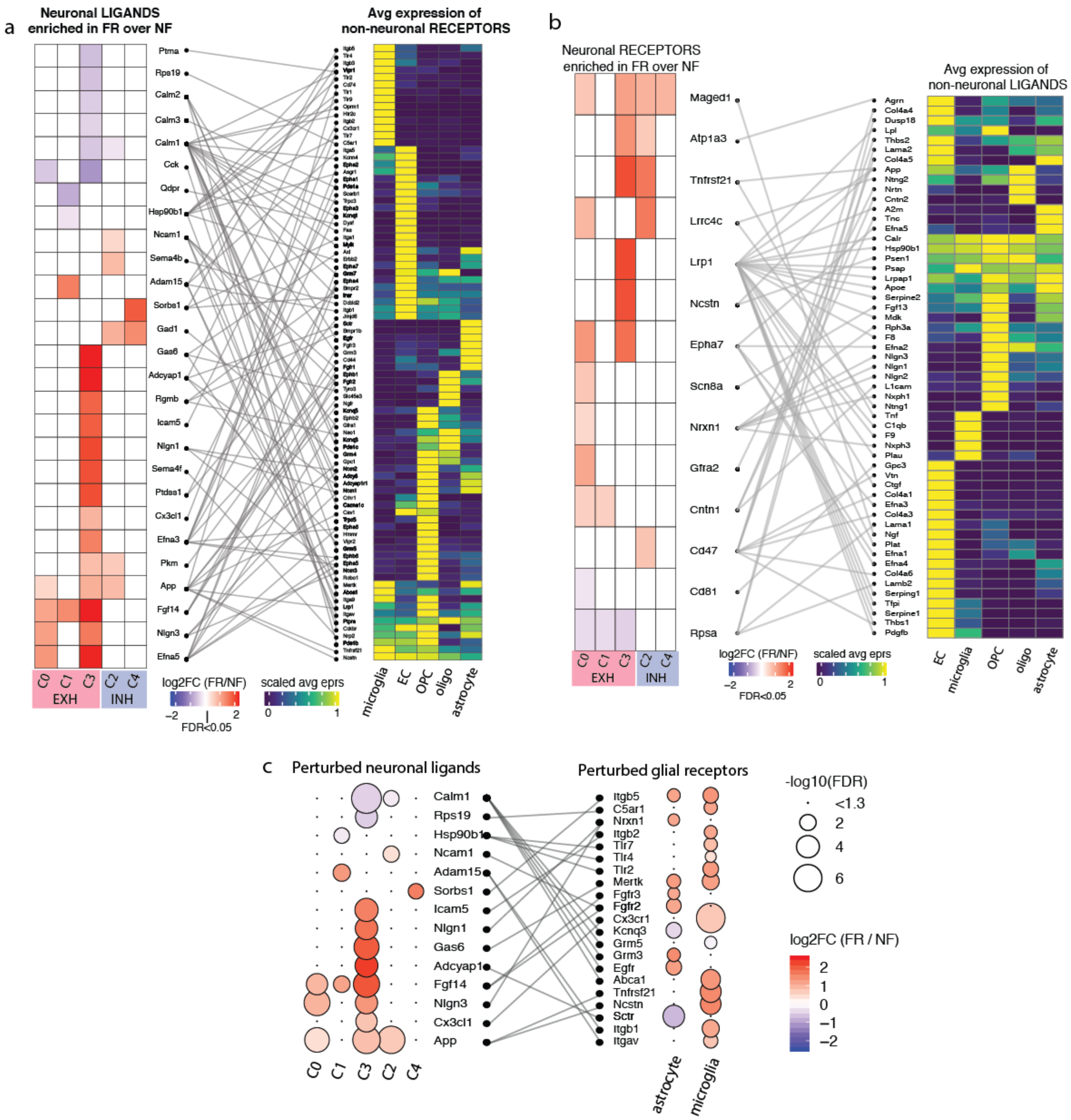


**Extended Fig 8 (related to Fig 5). Potential neuronal and non-neuronal cell ligand receptor-binding interactions**

a) (Left) Heatmap of the log2FC of DEGs (FR over NF) in neurons that are classified as ligands. (Middle and Right) Sankey plot of known ligand-receptor pairs (from Ramilowski et al, 2016) and heatmap of the average scaled expression level of the corresponding receptors in each type of non-neuronal cell.

b) (Left) Heatmap of the log2FC of DEGs (FR over NF) in neurons that are classified as receptors. (Middle and Right) Sankey plot of known ligand-receptor pairs and heatmap of the average scaled expression level of the corresponding ligands in each type of non-neuronal cell.

c) Dotted heatmap of a subset of neuronal ligands and glial receptors that are found to be differentially perturbed upon memory consolidation. Only receptors and ligands which were found to be (differentially) expressed are shown.

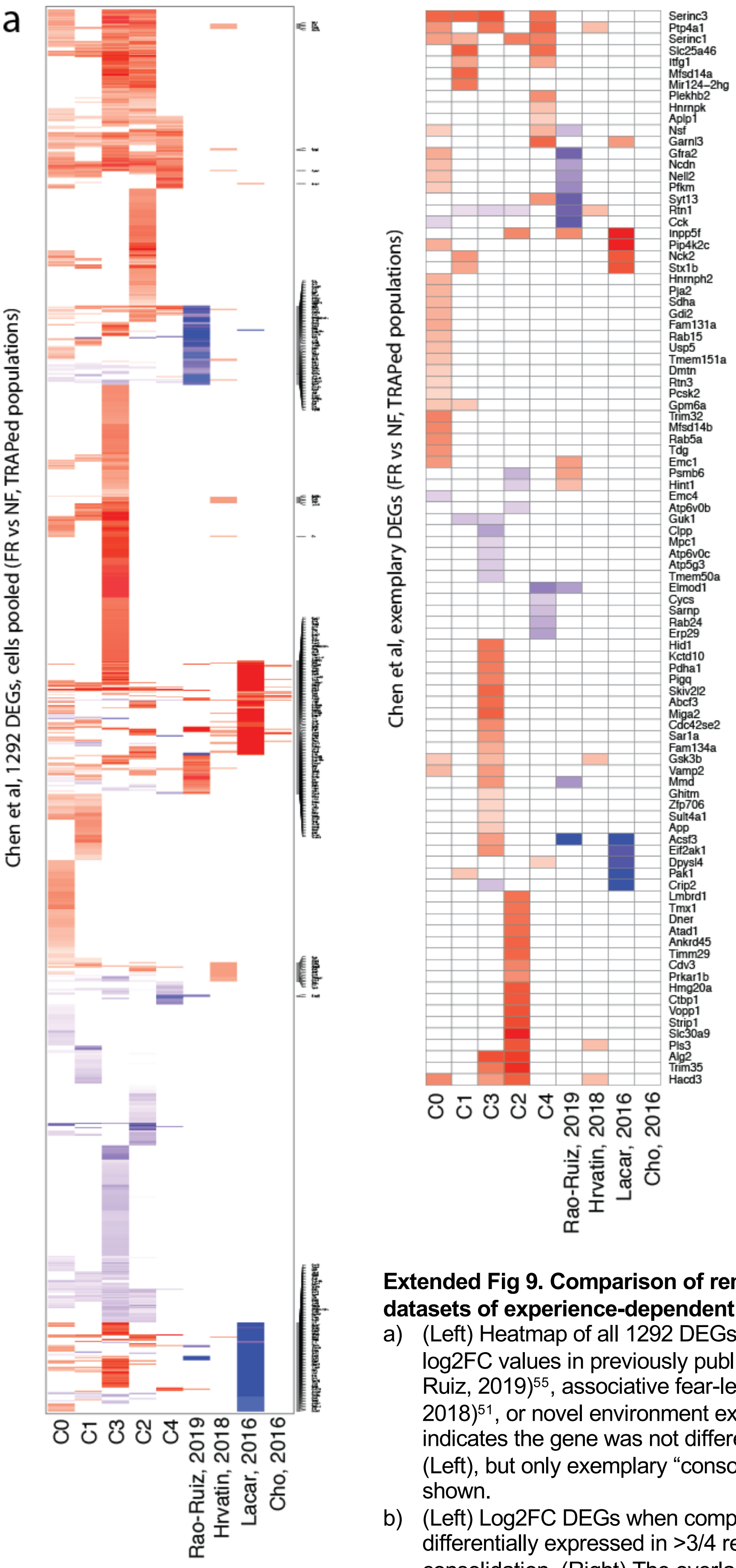


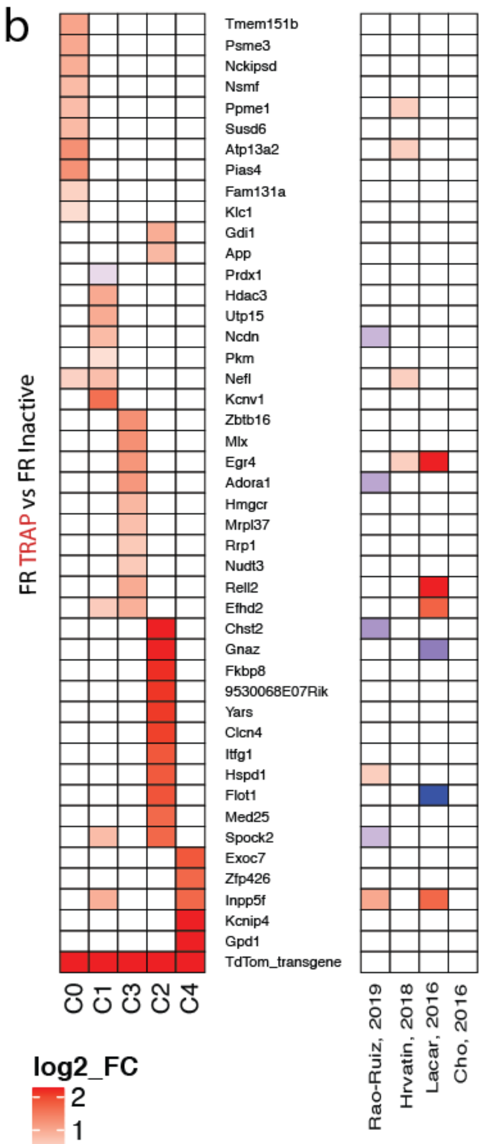


**Extended Fig 9. Comparison of remote memory DEGs with previously published datasets of experience-dependent transcriptional activity**

a) (Left) Heatmap of all 1292 DEGs (all cells pooled) in this manuscript, and their log2FC values in previously published datasets: recent fear memory retrieval (Rao-Ruiz, 2019)[55], associative fear-learning (Cho, 2016)[56], post-visual stimulus (Hvratin, 2018)[51], or novel environment exposure (Lacar, 2016)[57]. A value of zero log2FC indicates the gene was not differentially expressed in a dataset. (Right) same as (Left), but only exemplary "consolidation-dependent DEGs " from this study is shown.

b) (Left) Log2FC DEGs when comparing tdT+ vs tdT- neurons in FR mice (genes are differentially expressed in >3/4 replicates) undergoing remote fear memory consolidation. (Right) The overlap of differential expression in other datasets (in B).

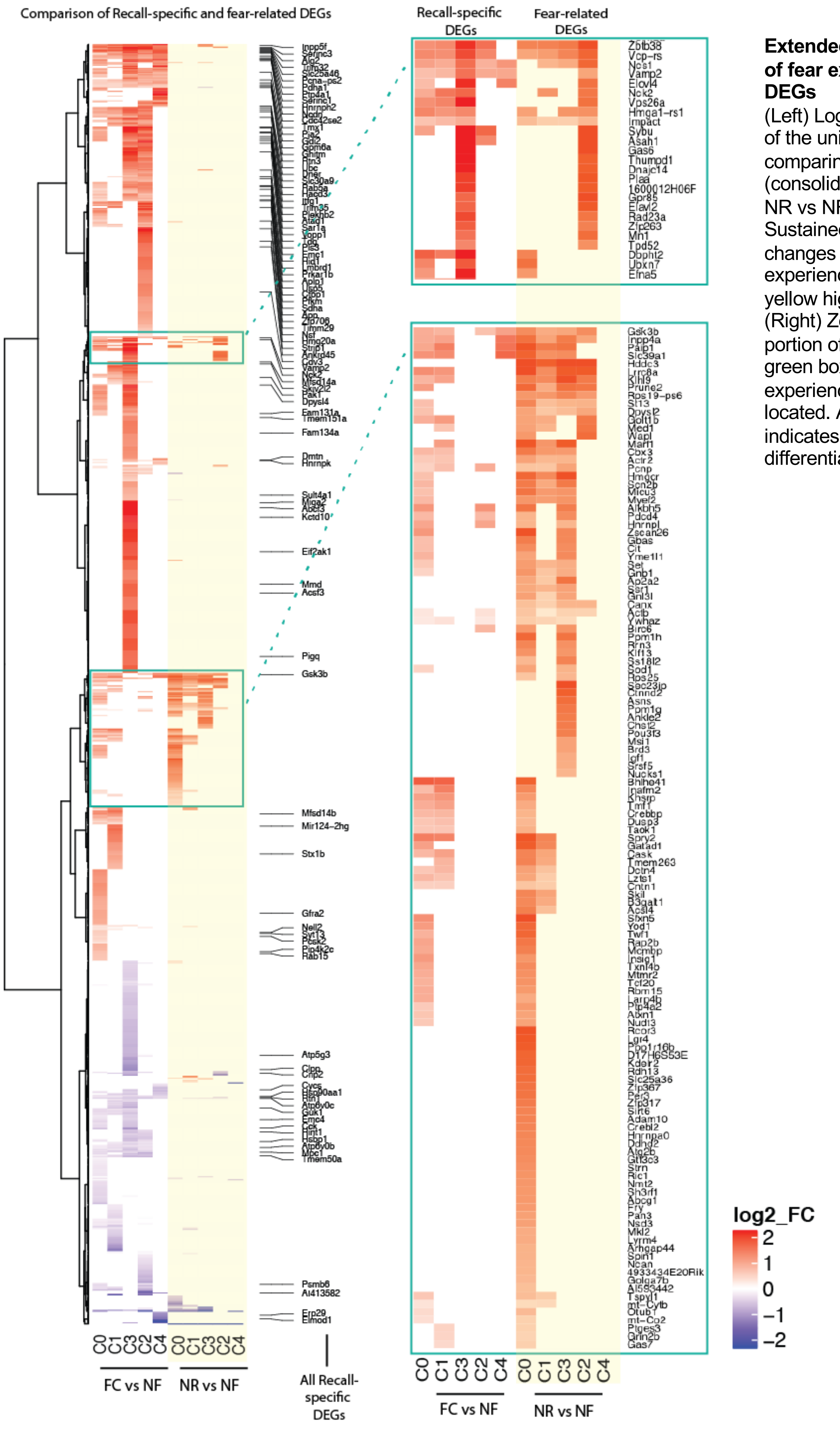


**Extended Fig 10. Identification of fear experience-related DEGs**

(Left) Log2 fold change heatmap of the union of DEGs from comparing FC vs NF (consolidation-dependent) and NR vs NF (Fear-related). Sustained transcriptional changes from the fear-experience itself is shown in the yellow highlighted columns. (Right) Zoomed in view of the portion of the heatmap within the green boxes where most fear experience-related DEGs are located. A log2FC of zero indicates that the gene is not differentially expressed.
